# Characterizing the Cancer-Associated Microbiome with Small RNA Sequencing Data

**DOI:** 10.1101/807586

**Authors:** Wei-Hao Lee, Kai-Pu Chen, Kai Wang, Hsuan-Cheng Huang, Hsueh-Fen Juan

**Author notes:** Corresponding author Hsuan-Cheng Huang, Institute of Biomedical Informatics, National Yang-Ming University, No.155, Sec.2, Linong Street, Taipei 112, Taiwan., Hsueh-Fen Juan, Department of Life Science, Institute of Molecular and Cellular Biology, Graduate Institute of Biomedical Electronics and Bioinformatics, National Taiwan University, No. 1, Sec. 4, Roosevelt Road, Taipei, 10617, Taiwan. **Wei-Hao Lee** is a research assistant in the Department of Life Science, National Taiwan University. His research interests include bioinformatics and cancer microbiome analysis. **Kai-Pu Chen** is a PhD student in the Graduate Institute of Biomedical Electronics and Bioinformatics, National Taiwan University. His research interests include network biology and cancer microbiome. **Dr. Kai Wang** is a principal scientist at Institute for Systems Biology, Seattle. His interest is studying the function, trafficking and application of extracellular RNAs. **Dr. Hsuan-Cheng Huang** is a professor in the Institute of Biomedical Informatics, National Yang-Ming University. His research interests include bioinformatics, computational and systems biology, and network biology. **Dr. Hsueh-Fen Juan** is a professor in the Department of Life Science, National Taiwan University. Her research interests include integrating omics and bioinformatics for biomarker and drug discovery.

## Abstract

The microbiome is recognized as a quasi-organ in the human body. In particular, the gut microbiome is correlated with immune function, metabolism, and tumorigenesis. When dysbiosis of the microbiome occurs, this variation may contribute to alterations in the microenvironment, potentially inducing an inflammatory immune response and providing a niche for neoplastic growth. However, there is limited evidence regarding the correlation and interaction between the microbiome and tumorigenesis. By utilizing microRNA sequencing data of patients with colon and rectal cancer from The Cancer Genome Atlas, we designed a novel analytical process to extract non-human small RNA sequences and align them with the microbial genome to obtain a comprehensive view of the cancer-associated microbiome. In the present study, we identified > 1000 genera among 630 colorectal samples and clustered these samples into three distinctive colorectal enterotypes. Each cluster has its own distinctive microbial composition and interactions. Furthermore, we found 12 genera from these clusters that are associated with cancer stages and revealed their putative functions. Our results indicate that the proposed analytical approach can effectively determine the cancer-associated microbiome. It may be readily applied to explore other types of cancer, in which specimens of the microbiome are difficult to collect.

## Introduction

The microbiome is a complex community in the human body, which comprises an immense number of microbes. It exhibits delicate interplay with the host activities, such as shaping the immune system, maintaining epithelium homeostasis, and regulating metabolism (1). The composition of the microbiome in various sites of the human body differs enormously. For example, the principal phyla in the colon and rectum are *Firmicutes* and *Bacteroidetes* (2), while the predominant phyla in breast tissue are *Proteobacteria* and *Firmicutes* (3). When the composition of the microbial community changed, relationship between microorganisms and host may also be altered, which may promote the development of different diseases. For example the involvement of altered microbiome have been reported in asthma (4), inflammatory bowel disease (5), and depression and anxiety (6).

The investigation of cancer-associated microbiome has attracted considerable attention which may lead towards the development of promising new targets in cancer diagnostics and therapeutics. Along with tumor growth, alterations in the composition of the microbiome may augment the severity of the disease and influence the tumor microenvironment (7,8). A recent study showed that some bacteria, for instance *Mycoplasma hyorhinis*, have the ability to modify the structure of gemcitabine to render its anticancer activity (9). Furthermore, microbes, such as *Fusobacterium nucleatum* reside adjacent to tumor may affect immune response and severity of cancer (10).

The 16S rDNA sequencing is the prevalent technique in studying bacteria, which provides researchers a rich source of information to distinguish a myriad of bacteria. The 16S rRNA gene is presented in all bacteria and its function and sequence are preserved through evolution (11). One of the challenges of using 16S rDNA is its data type which restricts the possibility of using large amount of existing data for example The Cancer Genome Atlas (TCGA) which provides various omics data of different types of cancers. In this report, we proposed a new analytical method for characterizing cancer-associated microbiome using the available small RNA-Seq data from cancer patients (Figure 1).

**Figure 1.**
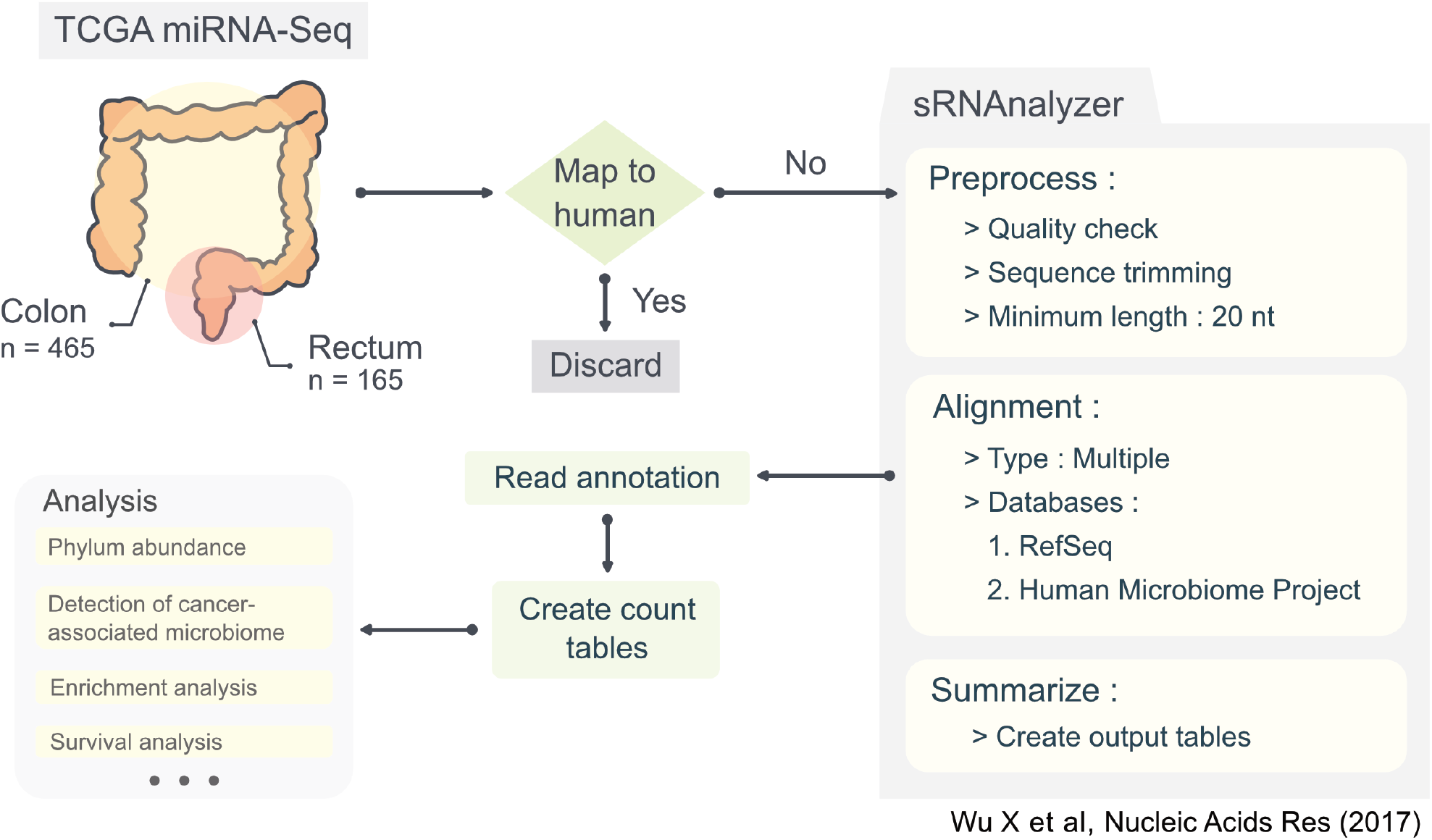
Flow chart of the analysis and detail information for miRNA-Seq data processing. This diagram illustrates the analysis procedures, including input data, tools used, and critical processes during analysis. The diagram also provides several crucial parameters which is required for the sRNAnalyzer pipeline to yield more accurate alignment results.

To examine whether our method has the same efficiency of identifying cancer-associated microbiome as 16S rDNA sequencing did on colorectal cancer (CRC) as well as the colon and rectum cancers, we took the CRC small RNA sequencing results from TCGA dataset as the subject of our study. We gathered CRC microRNA sequencing (miRNA-Seq) data from TCGA and applied our proposed method to identify the microbiome in each sample. We determined > 1,000 genera in 630 samples and obtained similar results as those obtained through 16S rDNA sequencing in previous investigations (8,12–14). The findings demonstrated that our method has a strong capacity to reveal the cancer-associated microbiome in CRC and may also be applied to different types of cancers.

## Materials and Methods

### Data preparation

CRC miRNA-Seq bam files were collected from the Genomic Data Commons Data Portal (downloaded: September 9, 2018). The files included two categories, namely colon adenocarcinoma (COAD; n = 465) and rectum adenocarcinoma (READ; n = 165) (15). The SAMtools (version 1.3.1) were used to extract small RNA reads, which were unmapped to the human genome in the bam files to generate possible reads originating from gut microbiota (16).

### Alignment of non-human small RNA reads

After filtering out the reads mapped to human miRNA and genome, we gathered non-human small RNA reads for microbiome analysis. The sRNAnalyzer pipeline was used for the alignment of non-human small RNA sequences (17). sRNAnalyzer is a specialized tool for alignment of small RNAs, which encompasses three procedures, namely “Preprocessing”, “Alignment”, and “Summarization”. Several parameters are required for data preprocessing and alignment. Accordingly, we set the minimum read length to 20 nucleotides, allowed reads to map to multiple reference databases for holistic, and omnibus mapping results, but not allowed any mismatch in order to obtain the highest accuracy of alignment. In the alignment procedure, we utilized the National Center for Biotechnology Information (NCBI) nucleotide database and Human Microbiome Project gastrointestinal database, provided through sRNAnalyzer, as our primary bacterial reference genome (17). However, we modified the procedure of the sRNAnalyzer after the alignment step and designed a new process customized for microbiome analysis.

### Annotation of bacterial small RNA reads

The alignment results was obtained with two Perl scripts (desProfile.pl and taxProfile.pl), which are provided in the sRNAnalyzer selfDev folder, to add information on reads including species taxonomy. The reads mapped to multiple species were assigned to their common taxon. Moreover, when the count table for each taxonomic rank was generated, reads from common taxonomic levels were assigned to a higher level. Therefore, those species-ambiguous reads could be assigned correctly to the right taxonomic ranks.

### Data Normalization

Since the microbiota count data contained a considerable number of zeros, we applied geometric mean of pairwise ratio (GMPR), a normalization method for zero-inflated data, to produce the count table (18). The GMPR normalization method requires two parameters, namely the intersection numbers for the minimum number of shared features and the minimum number of counts. The intersection numbers of the phylum, class, order, family, genus, and species count table were set at 1, 1, 3, 3, 5, and 5, respectively; the minimum number of counts we set was the default value provided by the package.

### Co-occurrence between genera

To visualize the bacterial genus co-occurrence network in each cluster, we calculated the relative genus abundance (percentage) in each sample, and selected genera that were detected in ≥ 20% of samples in the cluster. The Pearson correlation coefficient (PCC) was calculated and subsequently Fisher’s z-transformation was used to calibrate the PCC according to the sample size in each cluster. The pairs of genera with an absolute value of Fisher’s z-transformation ≤ 1.96 was removed. After removing uncorrelated pairs, Cytoscape was used for the computation of topological properties and network visualization (19).

### RNA-Seq expression data

CRC RNA expression data were downloaded from TCGA-COAD and TCGA-READ. Subsequently, the raw count was normalized with TMM normalization in the edgeR package (version 3.24.3) for cross-sample normalization (20).

### Functional gene set enrichment analysis

To demonstrate what biological processes of tumor cells are probably affected by cancer-associated bacteria, the Spearman correlation coefficient was calculated using the relative abundance of bacteria and patient RNA expression data. The fast pre-ranked gene set enrichment analysis (21) was applied to glean the candidate biological processes using the correlation coefficients as pre-ranked input data. The gene sets of biological processes in terms of Gene Ontology (GO) were downloaded from the Molecular Signatures Database (21, 22).

### Univariate survival analysis

The TCGA-COAD and TCGA-READ patient data were obtained using the R packages RTCGA (version 1.12.1) and RTCGA.clinical (version 20151101.12.0) (23). The survival analysis was conducted using cancer-associated bacteria as the univariate. We combined patient data with genus-relative abundance data, and categorized the samples into high and low sets using the median genus-relative abundance in 630 samples. This was performed using two R packages, namely survival (version 2.44-1.1) and survminer (version 0.4.3) (24, 25).

### Statistical analysis and data visualization

The R package vegan (version 2.5-4) was used for Shannon diversity and evenness analysis (26). Heatmaps were produced using the R package ComplexHeatmap (version 1.99.7) (27). Both the Kruskal–Wallis test and Mann–Whitney U test, shown as box plots, were performed using the R package ggpubr (version 0.2) (28). The networks was analyzed and computed their topological properties using Cytoscape (version 3.7.1) (19). Only the top 15 positive and negative correlations according to their Fisher Z-transformation value were depicted.

## Results

### Identification of CRC microbiota

To explore the microbiota associated with CRC (Figure 1), we mapped non-human small RNA reads against data from two microbial reference databases, namely the National Center of Biotechnology Information (NCBI) bacteria nucleotide and the National Institutes of Health (NIH) Human Microbiome Project. Approximately 626,084 unique small RNA sequences were aligned with those of microbial reference genomes, and ≤ 55.4% of the sequences aligned to specific genera or species. About 20% of the reads could be mapped to a myriad of bacteria from different phyla; thus, those sequences were excluded from further analysis. To ensure that the count data were not biased due to different library size in each sample, the raw count data was consequently calibrated using GMPR (18), a specialized normalization method for zero-inflated data like microbiome sequencing data, to produce the count table at each taxonomic rank.

### Composition of the microbiome in colorectal cancer

The relative abundance of phyla within the tumor and adjacent normal tissues was investigated to determine the phylogenetic composition of the microbiome in colorectal cancer (Figure 2A, B). The three predominant phyla identified were *Firmicutes*, *Bacteroidetes*, and *Proteobacteria*, which are the prevailing phyla in the human gut microbiota (29). The fourth and fifth most abundant phyla were *Actinobacteria*, and *Fusobacteria*, respectively. *Fusobacteria* were highly enriched in tumor tissues (Figure 2B), as shown by previous studies (10,30,31). *Fusobacterium nucleatum*, one of species in the *Fusobacteria* phylum, is usually detected in CRC and has the ability to generate a proinflammatory microenvironment (10, 30). In the current study, *Fusobacterium nucleatum* also had higher abundance in tumor samples (Supplementary Figure S1C). Furthermore, *Fusobacterium* was the third most abundant genus in 630 samples analyzed in the current study (Supplementary Figure S1A). Compared with previous research on the healthy gut microbiome (29), several of the reported genera were also found in our analysis. However, their relative abundance and the order of genera abundance were different. These findings demonstrated that several genera may correlate with CRC based on their abundance differences with the normal tissues.

**Figure 2.**
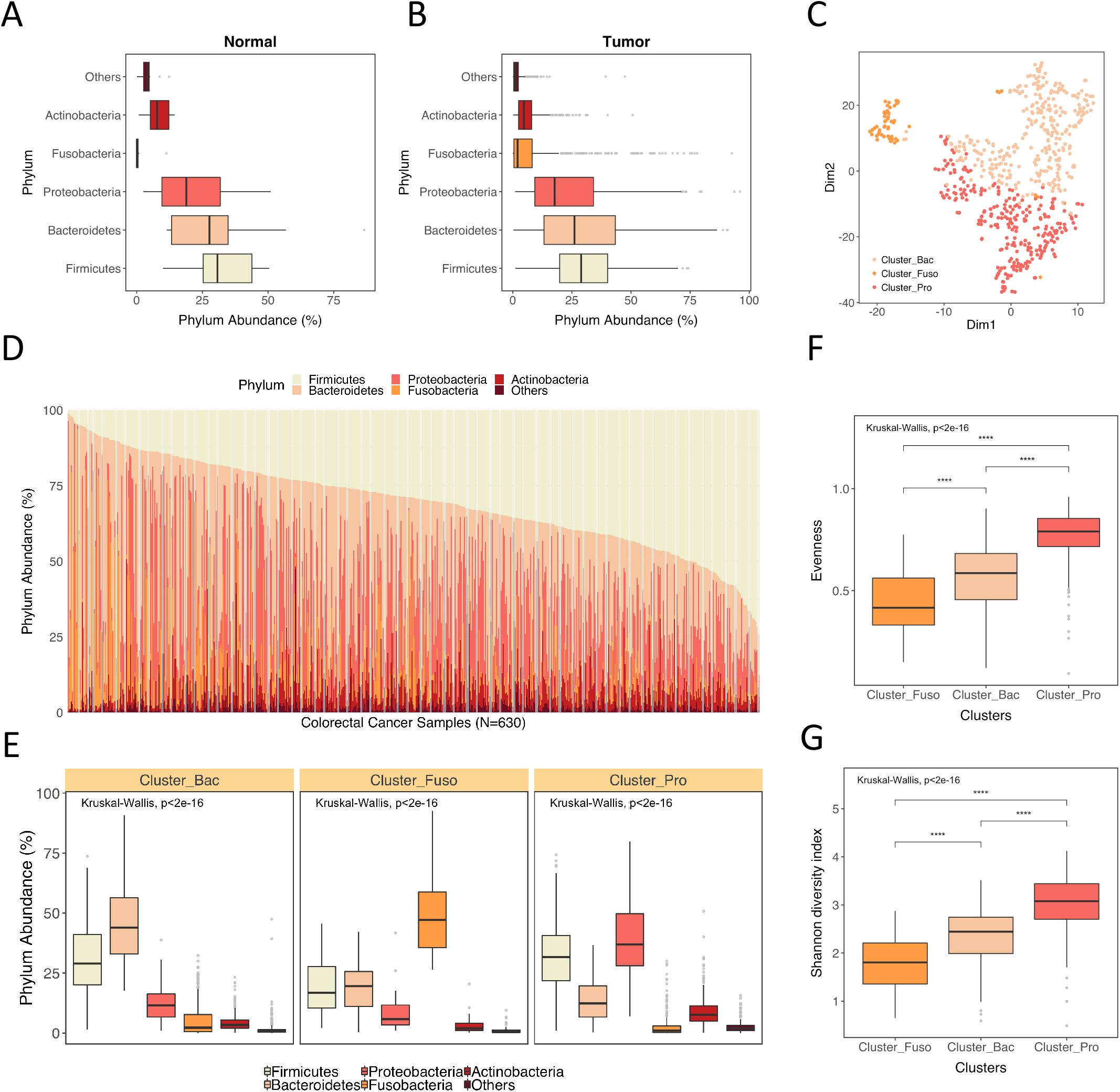
Phylogenetic profiles of the human gut microbiome in colorectal cancer. (A, B) Box plots of the relative abundance of bacterial phyla in solid normal tissues (N = 11) and tumor tissues (N = 616). (C) t-Distributed Stochastic Neighbor Embedding (tSNE) was used to perform spherical *k*-means cluster analysis. Colors represent clusters. (D) Stacked bar plot showing the five most abundant phyla in all samples and the rest of phyla grouped as “other”. (E) Box plots of phylum composition in the three different clusters, which were mostly composed of *Bacteroidetes* (Cluster_Bac and N = 306), *Fusobacteria* (Cluster_Fuso and N = 56), and *Proteobacteria* (Cluster_Pro and N = 268). (F), (G) Box plots showing diversity and evenness in three different clusters with significant difference. The Kruskal–Wallis test was used for comparison between multiple clusters and the Mann–Whitney U test was used for comparison of two clusters (shown with the star sign; (*p ≤ 0.05 and p > 0.01; **p ≤ 0.01 and p > 0.001; ***p ≤ 0.001).

The human gut microbiota is affected by diet, age, and other factors, resulting in a distinctive gut microbiota in each individual (32). The spectrum of bacterial phylum in all samples analyzed in the study is presented in Figure 2D and the results showed unique composition for each individual. The samples were then grouped into 3 clusters, Cluster_Bac, Cluster_Fuso, and Cluster_Pro, based on relative phylum abundance using spherical *k*-means. The box plots, t-distributed stochastic neighbor embedding (t-SNE; Figure 2C), as well as principle component analysis (Supplementary Figure S2) were used to illustrate microbiome composition differences among the three clusters. The names of these clusters were derived from the most abundant phylum identified in each cluster (Figure 2E; Supplementary Figure S3). To examine whether the clustering results were affected by the location of cancer (colon vs. rectum), the t-SNE results was replotted based on cancer location (Supplementary Figure S1B). TCGA-COAD and TCGA-READ were evenly distributed in the t-SNE plot, which suggested the clustering results were influenced by bacterial compositions rather than cancer location. We observed the differences in Shannon-diversity and evenness index between the three clusters (Figure 2F). Cluster_Pro exhibited the highest diversity and the most genera abundance distributed evenly in predominant genera compared to other clusters. Interestingly, the patients’ guts that was highly enriched with *Fusobacteria*, Cluster_Fuso showed the lowest diversity and evenness in their microbiomes. However, the high abundance of *Fusobacteria* did not show a strong correlation with cancer stage (Table 1).

**Table 1.** Information regarding the samples in TCGA-COAD, TCGA-READ, and the three clusters.

| *TCGA Colorectal Cancer miRNA* |  |  |  |  |  |  |
| --- | --- | --- | --- | --- | --- | --- |
|  |  | Original Data |  | After Clustering |  |  |
|  |  | TCGA COAD | TCGA READ | Cluster_Fuso | Cluster_Pro | Cluster_Bac |
| Sample Size | Solid Tissue Normal | 8 | 3 | 0 | 5 | 6 |
|  | Tumor | 457 | 162 | 56 | 263 | 300 |
| Gender | Male | 239 | 87 | 29 | 140 | 148 |
|  | Female | 224 | 77 | 27 | 126 | 157 |
| BMI | Mean / Median | 30.18 / 26.83 | 26.96 / 26.49 | 26.66 / 26.95 | 29.88 / 27.76 | 29.31 / 26.22 |
| Age at diagnosis | Mean / Median | 67.46 / 69 | 64.37 / 66 | 66.09 / 66.5 | 66.09 / 68 | 67.25 / 68 |
| Cancer Stage | Stage i | 75 | 29 | 10 | 52 | 42 |
|  | Stage ii | 175 | 48 | 22 | 78 | 123 |
|  | Stage iii | 129 | 50 | 17 | 82 | 80 |
|  | Stage iv | 65 | 24 | 7 | 34 | 48 |
|  | Not Reported | 13 | 11 | 0 | 17 | 7 |

Note :1. Tumor samples contain 2 recurrent tumor, 1 metastatic, and others are primary tumor.
2. The p-value of Pearson's Chi-squared test with different cluster and cancer stage (without not reported) is 0.1325.

Since the differences in the phylum composition observed in the three clusters, we constructed genus co-occurrence networks (Figure 3A–C) and heatmaps (Supplementary Figure S4A–C). We further determined the topological properties of each genus to gather more evidence regarding the interplays of genera (Supplementary Table S1–3). Due to the complex correlation in the full co-occurrence network, only the top 15 positive and negative correlations based on their Fisher Z-transformation values were showed in Figure 3. These networks illustrated that each sample cluster exhibited distinctive bacterial interplay and different hub genera. For instance, *Bacteroides* had the third highest network degree in the Cluster_Bac co-occurrence network (Supplementary Table S1, Figure 3A), and the highest betweenness centrality. This indicated that *Bacteroides* played an important role that might affect other genera and acted as a hub in this network. In the Cluster_Fuso co-occurrence network (Supplementary Table S2, Figure 3B), although *Fusobacterium* did not have the highest network degree, *Fusobacterium* had several strong correlations in the network, indicating that *Fusobacterium* is also a hub in the co-occurrence network.

**Figure 3.**
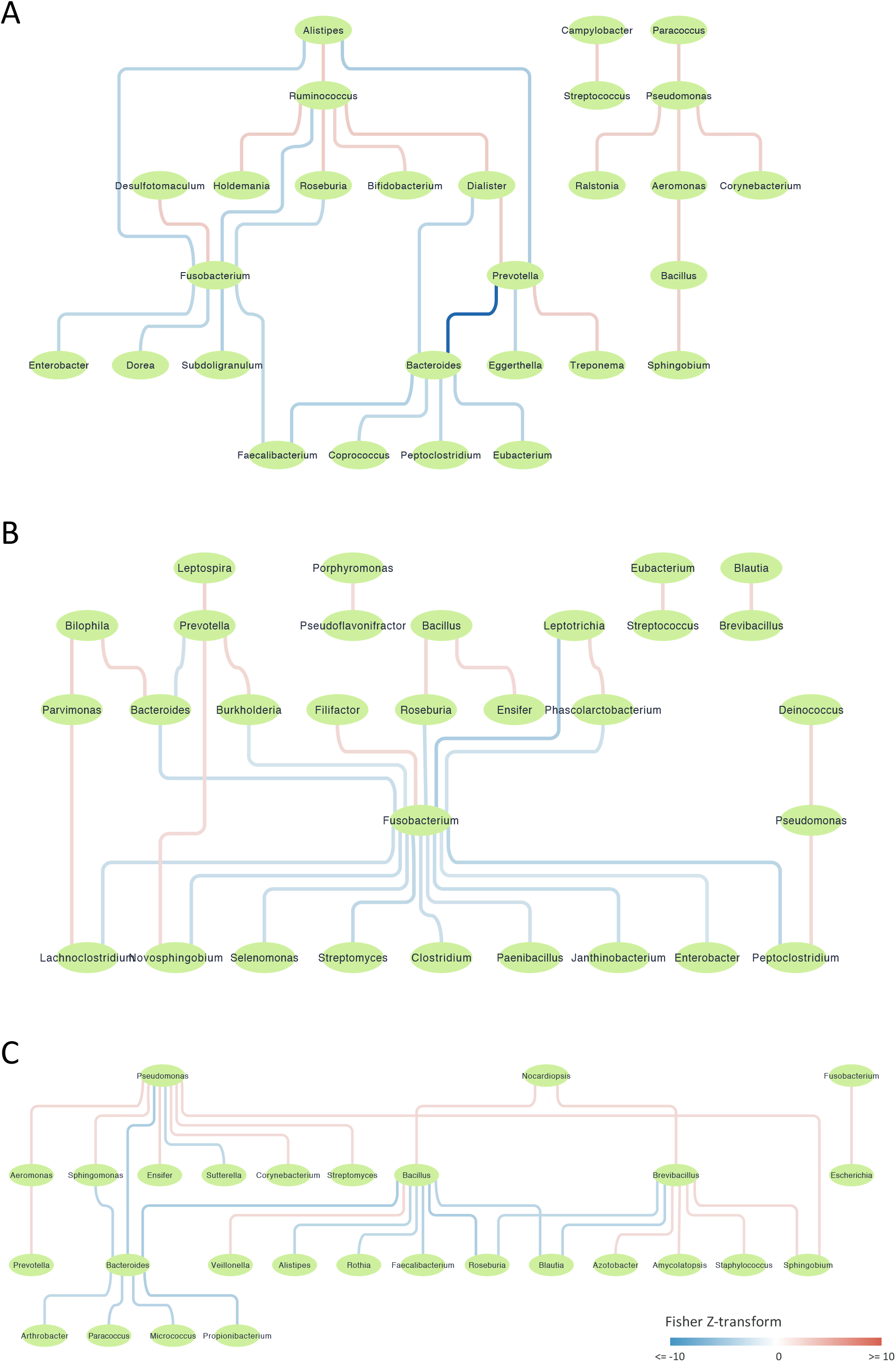
Bacterial co-occurrence networks in the three clusters. (A–C) Co-occurrence networks of bacterial genera in different clusters, namely (A) Cluster_Bac, (B) Cluster_Fuso, and (C) Cluster_Pro. The correlations of edges were calculated using the Pearson correlation coefficient. Fisher z-transformation was used for normalization according to sample size, rendering the networks comparable. We only noted the top 15 positive correlations and top 15 negative correlations in these three networks.

### Detection of cancer-associated bacteria

To detect cancer-associated bacteria, we first selected common bacterial genera which were detected in ≥ 20% of the samples. The genus-relative abundance in normal tissues adjacent to tumors in the three clusters were compared and calculated. More than 50 genera with differential abundance in cancer were observed (Supplementary Figure S5).

The 25 genera previously reported as cancer-associated bacteria were used to examine the consistency of our results with other studies (Figure 4) (7,8,12–14,31,33–37). Several genera, such as *Porphyromonas, Roseburia, Ruminococcus, Subdoligranulum, Bacterodies*, and *Prevotella*, were reportedly increased as well as decreased in CRC; hence, we classified these genera in an ambiguous category.

**Figure 4.**
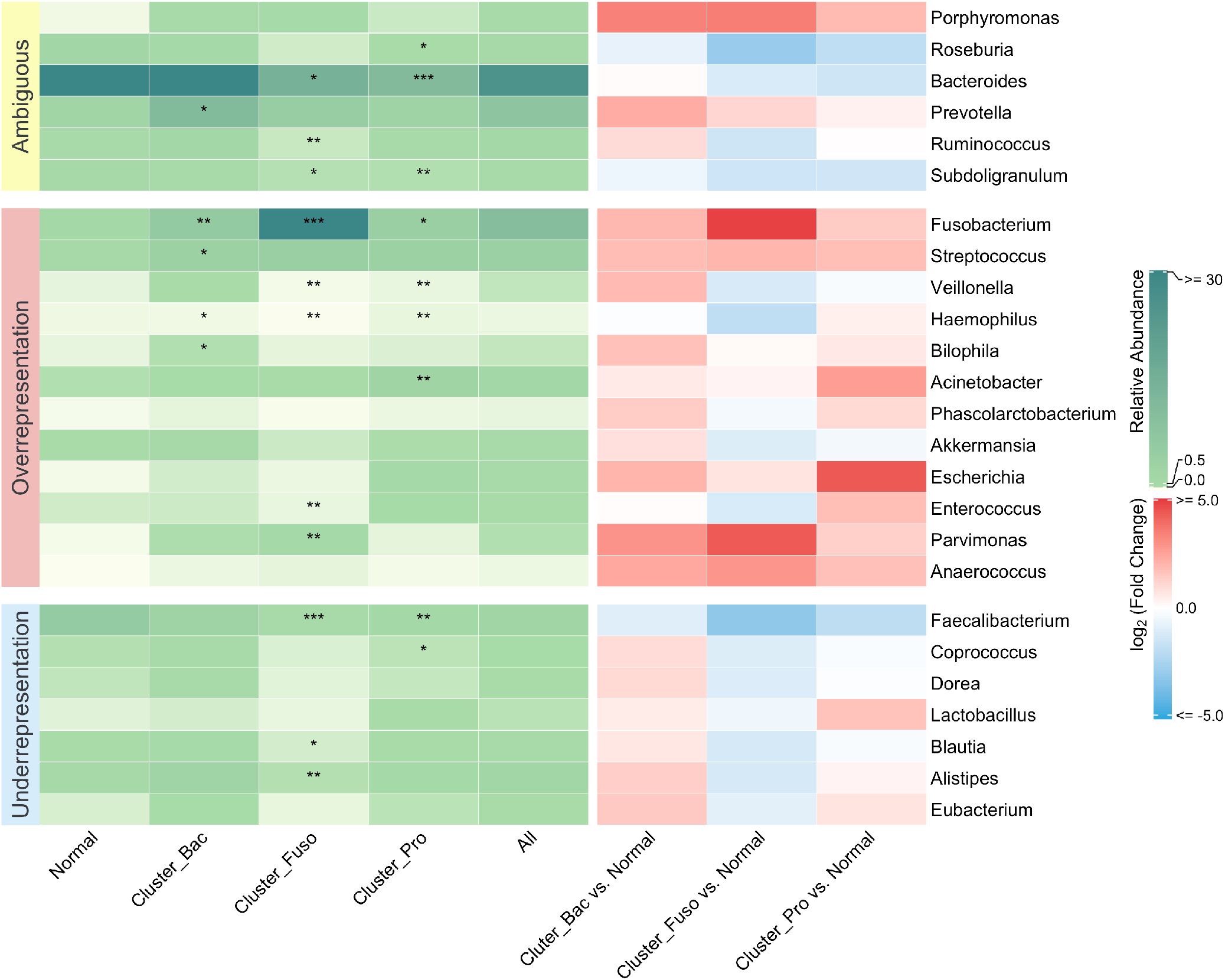
Cancer-associated microbiome consistent with previous studies. Previous studies reported 25 genera as the cancer-associated microbiome. We categorized those genera into three groups based on their representation in CRC: ambiguous, over-representation, and under-representation. Genera in the ambiguous category were found to be both overrepresented and underrepresented in previous studies. The left heatmap (colored green) indicates the relative abundance of genera, and each column stands for three clusters (normal adjacent samples were removed), normal adjacent samples, and all samples. The statistical significance of the three clusters was calculated between tumors in each cluster and all normal adjacent tissues using the Mann–Whitney U test (*p ≤ 0.05 and p > 0.01; **p ≤ 0.01 and p > 0.001; ***p ≤ 0.001). The right heatmap colored with log2 (fold change). Fold change was defined as the genus relative abundance in clusters divided by the genus relative abundance in normal samples.

In the category of over-represented bacteria in the cancer-associated microbiome, the abundance of *Fusobacterium*, *Streptococcus*, *Veillonella*, *Haemopilus*, *Bilophila*, *Acinetobacter*, *Phascolarctobacterium*, *Akkermansia*, *Escherichia*, *Enterococcus*, *Parvimonas*, and *Anaerococcus* was consistent with previous studies (12,14,33,34,37). However, in Cluster_Fuso these showed lower abundance compared to normal samples. In the under-represented category, a portion of the genera in Cluster_Bac and Cluster_Pro showed higher abundance than normal samples. According to the complete patient clinical data, the correlations between bacterial genus abundance and different cancer stages were observed. Both *Dorea* and *Blautia* showed a tendency toward decreasing abundance along with an advanced cancer stage, despite having a higher average abundance than normal samples. Twelve genera including *Dorea*, *Faecalibacterium*, *Roseburia*, *Pseudoflavonifractor*, *Sutterella*, *Pseudomonas*, *Blautia*, *Faecalitalea*, *Bacteroides*, *Subdoligranulum*, *Amycolatopsis*, and *Desulfovibrio* were considered cancer stage-associated bacteria in different clusters (Figure 5). Most of the cancer stage-associated genera, such as *Dorea*, *Roseburia*, and *Blautia*, exhibited a tendency of decreasing abundance along with advanced cancer stage. Only three genera, *Faecalitalea*, *Amycolatopsis*, and *Desulfovibrio*, showed higher abundance in stage IV cancer.

**Figure 5.**
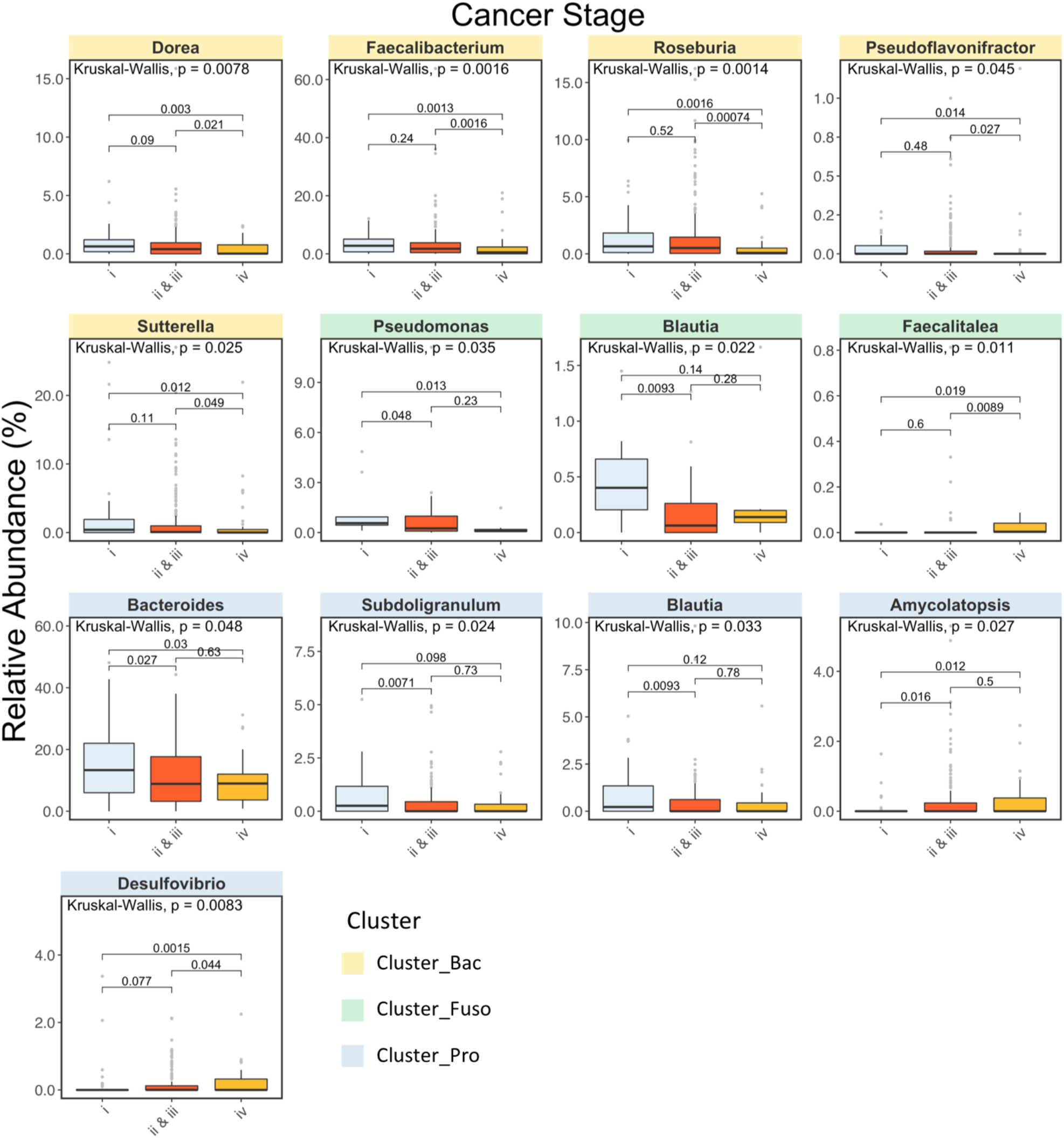
Cancer stage-associated microbiome in the three clusters. Twelve genera are cancer stage-associated bacteria, and each box plot shows one genus. The facet color indicates the cluster in which a genus was identified. Stages shown on the x axis were clustered into three categories (stage I, stage II & III, and stage IV). P-values labeled in the box plot using the Kruskal–Wallis test for comparison between multiple stages and the Mann–Whitney U test for comparison between two stages (*p ≤ 0.05 and p > 0.01; **p ≤ 0.01 and p > 0.001; ***p ≤ 0.001).

### Biological functions correlate with cancer-associated bacteria

The finding of cancer stage-associated microbiome population allows us to explore the relationships between bacterial abundance and tumor progression, and the possible influence of microbiome on host in terms of the perturbed biological processes. The correlation between bacterial genera abundance and mRNA gene expression was calculated using the Spearman correlation coefficient, and applied pre-ranked gene set enrichment analysis methods to identify the enriched biological processes influenced by cancer stage-associated bacteria. The top 50 most representative enriched biological processes (in GO terms) were selected and displayed their enrichments in different cancer-associated bacteria in Figure 6. Among these functions, immune responses (i.e., leukocyte activation, leukocyte degranulation, regulation of the T-cell receptor signaling pathway, etc.) were highly correlated with the cancer stage-associated microbiome. Moreover, several metabolic processes (i.e., fatty acid and carbohydrate catabolic processes, oligosaccharide metabolic process, etc.) were also related to the cancer-associated microbiome. These results provided a holistic picture of plausible effects caused by the microbiome and need to be scrutinized by additional studies.

**Figure 6.**
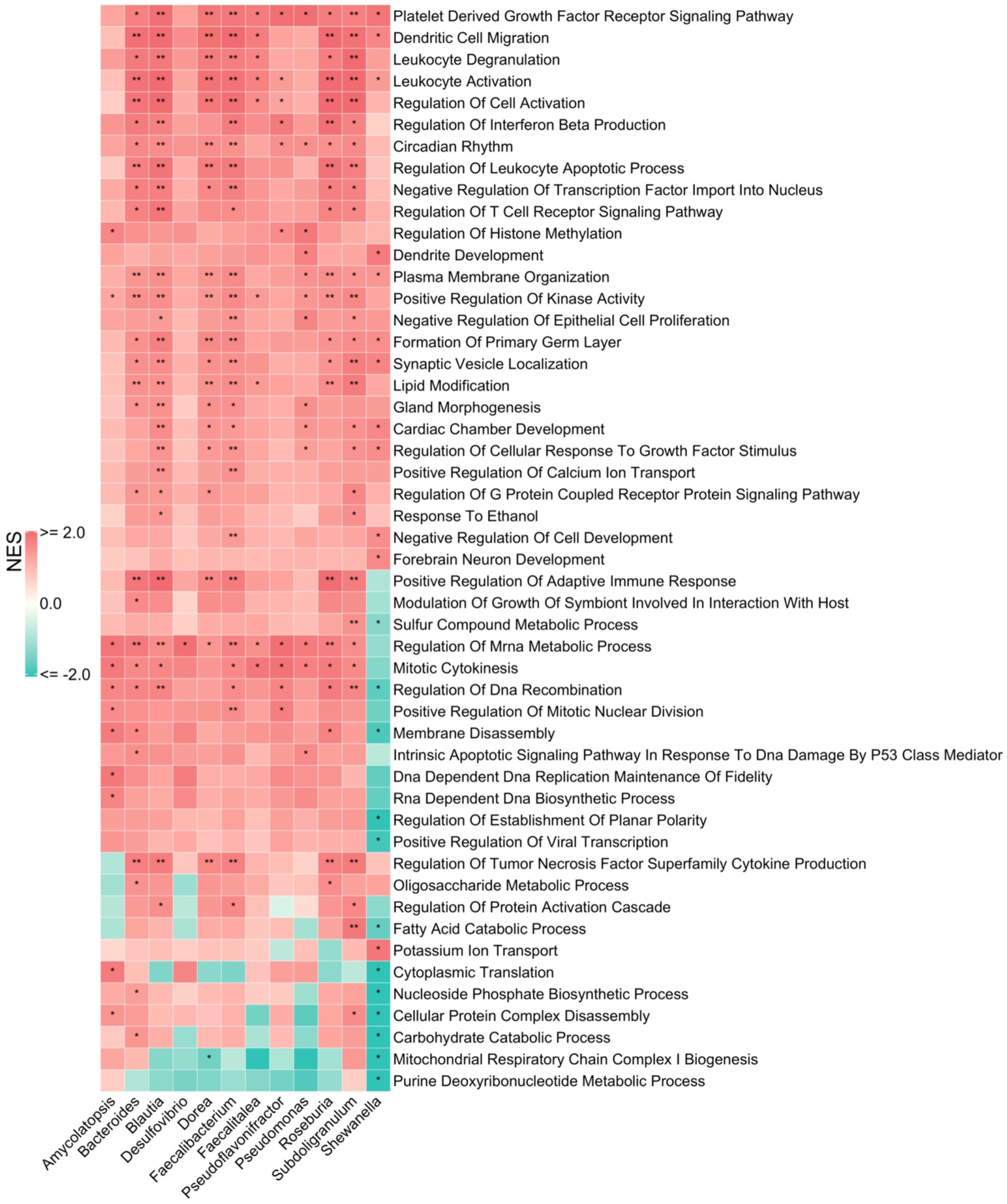
Gene ontology related to the cancer stage-associated microbiome. We calculated the correlation between biological processes in gene ontology and the cancer stage-associated microbiome using gene set enrichment analysis. The heatmap presents 50 enriched biological processes and the color indicates normalized enrichment score (NES). *adjusted p ≤ 0.05 and p > 0.01; **adjusted p ≤ 0.01 and p > 0.001;***adjusted p ≤ 0.001.

### Correlation between bacteria and patient survival

Survival analysis was conducted to explore whether cancer stage-associated bacteria were correlated with patient survival. The patients were into high- and low-abundance groups according to the cancer stage-associated microbiome, and found that four genera (*Dorea*, *Blautia*, *Subdoligranulum*, and *Sutterella*) had a strong correlation with patient survival. Higher relative abundance of those bacteria was associated with better survival (Figure 7). A previous study showed that *Blautia* in the gut microbiome might reduce mortality from graft-versus-host disease (38). Therefore, these four genera may provide new targets for the treatment of patients with poor prognosis.

**Figure 7.**
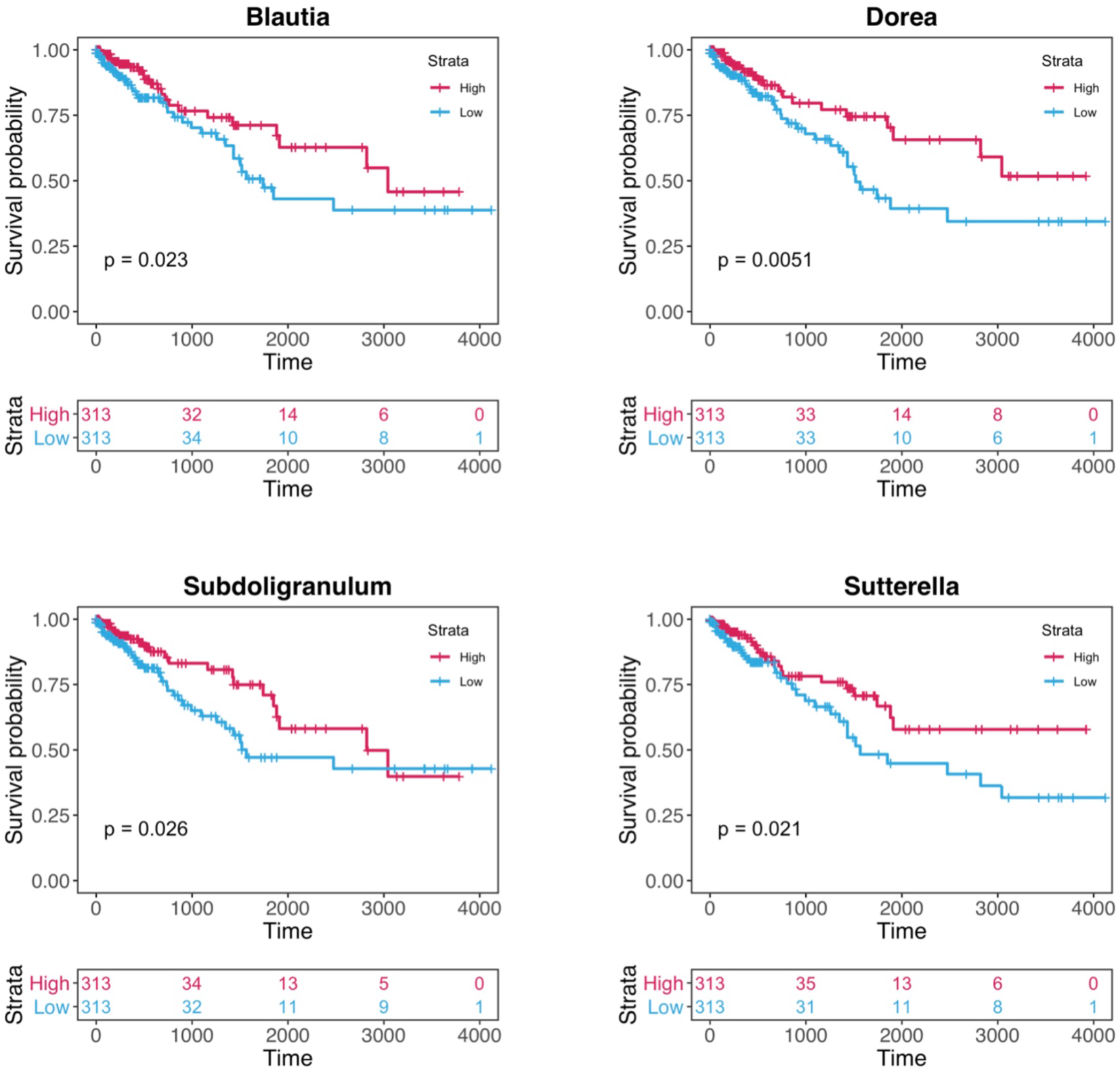
Four cancer stage-associated genera correlated with patient survival. Kaplan–Meier plots illustrate that the relative abundance of four genera correlates with survival. Strata below the Kaplan–Meier plot indicates the number of patients in the high-and low-abundance groups.

## Discussion

A novel analytical approach was developed and used to demonstrate the possibility of using the small RNA sequencing data to study cancer-associated microbiome. This approach provided results, which were similar to those reported in previous studies using 16s rDNA sequencing (7,8,12–14,31,33–37). We utilized non-human miRNA reads extracted from TCGA-COAD and TCGA-READ miRNA-Seq data. These miRNA-Seq data were directly collected from tumor and adjacent normal tissues (39). After extracting non-human small RNA reads, we performed sequence alignment against various bacterial sequence databases with multiple stringent parameters to yield more accurate results, and filtered out unqualified reads during alignment. Owing to the fact that short reads may be mapped to multiple species, we considered all possible results and annotated those reads with their joint taxonomic rank. In addition, approximately half of the reads that mapped to several phyla were excluded, and the remaining reads which could be identified in certain taxonomic ranks except phylum. This step is similar to the construction of operational taxonomic unit clusters using sequence similarity in 16s rDNA sequencing analysis (40).

The 16s rDNA sequencing is based on DNA so each bacterium has one copy; however, small RNA sequencing is based on RNA and each bacterium will have many copies. When growth condition is better, the levels of rRNA and mRNA in bacteria can be even higher. In theory, the RNA-based measurement can provide better picture of bacteria composition due to amplification of the signal from one copy (DNA) to thousands or millions copies (RNA); however, the amplification process may skew the composition since it favors the bacteria that are growing. Nevertheless, the results showed some similarities between the two platforms, suggesting that the dominate signal (composition) is faithfully “translated” from DNA to RNA but for weaker signals, the RNA might work better. In addition, the RNA-based measurement may also reflect the activity of the bacteria community — the less representing groups (in RNA-based measurement compared to DNA-based measurement) may be less active.

The five most abundant phyla in both tumor and adjacent normal tissues identified in this study (Figure 2A, B) were in agreement with previous findings (7,10,30,31). One of phyla, *Fusobacteria*, was highly enriched in tumor tissues (7,10,30,31). The results indicated that *Fusobacteria* prefer to inhabit tumor tissues rather than adjacent normal tissues (10). Moreover, that the results showed that each patient had a distinctive microbiome composition (Figure 2D) in tumor tissues, which might be affected by diet, drug treatment, age and disease stage. The differences of microbiome composition might produce unique microenvironments which correlate with host biological functions (1). Based on the spectrum of microbiome, the samples can be classified into three major clusters, namely Cluster_Bac, Cluster_Fuso, and Cluster_Pro (Figure 2E) using spherical *k*-means method. The adjacent normal tissues were collected from sites near tumors; therefore, those tissues would have the parallel dominant microbes as the tumor tissues. However, there is a small portion of microbes changed their abundance in tumor as well as different tumor stages, which probably caused by the inter-species competition in the tumor microenvironment (Figure 4-5; Supplementary Figure S5). The findings of cancer-associated microbiome through mining small RAN sequencing data are similar to prior reports (Figure 4) (7,8,12–14,31,33–37). Interestingly the genera in Cluster_Fuso exhibited different pattern from previous studies (8,13,33,41,42). *Fusobacteria* were highly abundant in Cluster_Fuso, and the genus *Fusobacterium* negatively correlated with other genera (Figure 3 and Supplementary Figure S3B). Therefore, genus representation in Cluster_Fuso may differ from comparing with that reported in previous studies (Figure 4) (7,8,12–14,31,33–37). *Dorea* and *Blautia* had higher abundance in Cluster_Bac and Cluster_Pro and the abundance of them declined in parallel with advancing cancer stage (Figure 5). These results were also consistent with previous studies (8,12,33,41,42).

Based on large-scale cohort studies and meta-omics resources from the TCGA, several techniques were applied to examine the correlation between the cancer-associated microbiome and biological functions in tumors. There is one caution of using small RNA sequencing data for microbiome analysis, which is short read length. Due to their short length, an immense amount of reads could be aligned to multiple species, and only part of the reads could be assigned to unique species. This limited the taxonomic resolution of the analysis. Nevertheless, in the taxonomic rank of genera, genus count data could provide a sufficient amount of reads for further analysis steps, and higher taxonomic ranks may yield more robust results.

In conclusion, this method provides a simple and robust approach for analyzing the cancer-associated microbiome based on small RNA sequencing data and the findings are similar to those obtained using the 16S rDNA method. Currently, colonization of numerous body sites by the microbiome has been reported, and an increasing number of studies are being conducted to investigate the association between cancer and the microbiome. Our method can be applied to different types of cancers and utilized results from large-scale cohort studies (e.g., using TCGA data) to identify links between patient variables and microbiome.

## Supplementary Data

Supplementary Data are available online.

## References

1. Belkaid, Y. and Hand, T.W. (2014) Role of the microbiota in immunity and inflammation. Cell, 157, 121–141.

2. Rinninella, E., Raoul, P., Cintoni, M., Franceschi, F., Miggiano, G.A.D., Gasbarrini, A. and Mele, M.C. (2019) What is the Healthy Gut Microbiota Composition? A Changing Ecosystem across Age, Environment, Diet, and Diseases. Microorganisms, 7.

3. Urbaniak, C., Gloor, G.B., Brackstone, M., Scott, L., Tangney, M. and Reid, G. (2016) The Microbiota of Breast Tissue and Its Association with Breast Cancer. Appl Environ Microbiol, 82, 5039–5048.

4. Sokolowska, M., Frei, R., Lunjani, N., Akdis, C.A. and O'Mahony, L. (2018) Microbiome and asthma. Asthma Res Pract, 4, 1.

5. Halfvarson, J., Brislawn, C.J., Lamendella, R., Vazquez-Baeza, Y., Walters, W.A., Bramer, L.M., D'Amato, M., Bonfiglio,F., McDonald, D., Gonzalez, A. et al. (2017) Dynamics of the human gut microbiome in inflammatory bowel disease. Nat Microbiol, 2, 17004.

6. Foster, J.A. and McVey Neufeld, K.A. (2013) Gut-brain axis: how the microbiome influences anxiety and depression. Trends Neurosci, 36, 305–312.

7. Ahn, J., Sinha, R., Pei, Z., Dominianni, C., Wu, J., Shi, J., Goedert, J.J., Hayes, R.B. and Yang, L. (2013) Human gut microbiome and risk for colorectal cancer. J Natl Cancer Inst, 105, 1907–1911.

8. Gao, Z., Guo, B., Gao, R., Zhu, Q. and Qin, H. (2015) Microbiota disbiosis is associated with colorectal cancer. Front Microbiol, 6, 20.

9. Geller, L.T., Barzily-Rokni, M., Danino, T., Jonas, O.H., Shental, N., Nejman, D., Gavert, N., Zwang, Y., Cooper, Z.A., Shee, K. et al. (2017) Potential role of intratumor bacteria in mediating tumor resistance to the chemotherapeutic drug gemcitabine. Science, 357, 1156–1160.

10. Kostic, A.D., Chun, E., Robertson, L., Glickman, J.N., Gallini, C.A., Michaud, M., Clancy, T.E., Chung, D.C., Lochhead, P., Hold, G.L. et al. (2013) Fusobacterium nucleatum potentiates intestinal tumorigenesis and modulates the tumor-immune microenvironment. Cell Host Microbe, 14, 207–215.

11. Janda, J.M. and Abbott, S.L. (2007) 16S rRNA gene sequencing for bacterial identification in the diagnostic laboratory: pluses, perils, and pitfalls. Journal of clinical microbiology, 45, 2761–2764.

12. Chen, W., Liu, F., Ling, Z., Tong, X. and Xiang, C. (2012) Human intestinal lumen and mucosa-associated microbiota in patients with colorectal cancer. PLoS One, 7, e39743.

13. Sun, J. and Kato, I. (2016) Gut microbiota, inflammation and colorectal cancer. Genes Dis, 3, 130–143.

14. Weir, T.L., Manter, D.K., Sheflin, A.M., Barnett, B.A., Heuberger, A.L. and Ryan, E.P. (2013) Stool microbiome and metabolome differences between colorectal cancer patients and healthy adults. PloS one, 8, e70803.

15. Grossman, R.L., Heath, A.P., Ferretti, V., Varmus, H.E., Lowy, D.R., Kibbe, W.A. and Staudt, L.M. (2016) Toward a Shared Vision for Cancer Genomic Data. N Engl J Med, 375, 1109–1112.

16. Li, H., Handsaker, B., Wysoker, A., Fennell, T., Ruan, J., Homer, N., Marth, G., Abecasis, G., Durbin, R. and Genome Project Data Processing, S. (2009) The Sequence Alignment/Map format and SAMtools. Bioinformatics, 25, 2078–2079.

17. Wu, X., Kim, T.K., Baxter, D., Scherler, K., Gordon, A., Fong, O., Etheridge, A., Galas, D.J. and Wang, K. (2017) sRNAnalyzer-a flexible and customizable small RNA sequencing data analysis pipeline. Nucleic Acids Res, 45, 12140–12151.

18. Chen, L., Reeve, J., Zhang, L., Huang, S., Wang, X. and Chen, J. (2018) GMPR: A robust normalization method for zero-inflated count data with application to microbiome sequencing data. PeerJ, 6, e4600.

19. Shannon, P., Markiel, A., Ozier, O., Baliga, N.S., Wang, J.T., Ramage, D., Amin, N., Schwikowski, B. and Ideker, T. (2003) Cytoscape: a software environment for integrated models of biomolecular interaction networks. Genome research, 13, 2498–2504.

20. Robinson, M.D., McCarthy, D.J. and Smyth, G.K. (2010) edgeR: a Bioconductor package for differential expression analysis of digital gene expression data. Bioinformatics, 26, 139–140.

21. Sergushichev, A. (2016) An algorithm for fast preranked gene set enrichment analysis using cumulative statistic calculation. BioRxiv, 060012.

22. Subramanian, A., Tamayo, P., Mootha, V.K., Mukherjee, S., Ebert, B.L., Gillette, M.A., Paulovich, A., Pomeroy, S.L., Golub, T.R. and Lander, E.S. (2005) Gene set enrichment analysis: a knowledge-based approach for interpreting genome-wide expression profiles. Proceedings of the National Academy of Sciences, 102, 15545–15550.

23. Kosinski, M. and Biecek, P. (2016) RTCGA: The cancer genome atlas data integration. R package version, 1.

24. Therneau, T.M. and Grambsch, P.M. (2013) Modeling survival data: extending the Cox model. Springer Science & Business Media.

25. Kassambara, A., Kosinski, M. and Biecek, P. (2017) survminer: Drawing Survival Curves using ‘ggplot2’. R package version 0.3, 1.

26. Oksanen, J. and Blanchet, F.G. Package ‘vegan’.

27. Gu, Z., Eils, R. and Schlesner, M. (2016) Complex heatmaps reveal patterns and correlations in multidimensional genomic data. Bioinformatics, 32, 2847–2849.

28. Kassambara, A. (2017) ggpubr:“ggplot2” based publication ready plots. R package version 0.1, 6.

29. Arumugam, M., Raes, J., Pelletier, E., Le Paslier, D., Yamada, T., Mende, D.R., Fernandes, G.R., Tap, J., Bruls, T., Batto, J.-M. et al. (2011) Enterotypes of the human gut microbiome. Nature, 473, 174–180.

30. Castellarin, M., Warren, R.L., Freeman, J.D., Dreolini, L., Krzywinski, M., Strauss, J., Barnes, R., Watson, P., Allen-Vercoe, E., Moore, R.A. et al. (2011) Fusobacterium nucleatum infection is prevalent in human colorectal carcinoma. Genome Research, 22, 299–306.

31. Kostic, A.D., Gevers, D., Pedamallu, C.S., Michaud, M., Duke, F., Earl, A.M., Ojesina, A.I., Jung, J., Bass, A.J., Tabernero, J. et al. (2012) Genomic analysis identifies association of Fusobacterium with colorectal carcinoma. Genome Res, 22, 292–298.

32. Yatsunenko, T., Rey, F.E., Manary, M.J., Trehan, I., Dominguez-Bello, M.G., Contreras, M., Magris, M., Hidalgo, G., Baldassano, R.N., Anokhin, A.P. et al. (2012) Human gut microbiome viewed across age and geography. Nature, 486, 222–227.

33. Wang, T., Cai, G., Qiu, Y., Fei, N., Zhang, M., Pang, X., Jia, W., Cai, S. and Zhao, L. (2011) Structural segregation of gut microbiota between colorectal cancer patients and healthy volunteers. The ISME Journal, 6, 320–329.

34. Geng, J., Song, Q., Tang, X., Liang, X., Fan, H., Peng, H., Guo, Q. and Zhang, Z. (2014) Co-occurrence of driver and passenger bacteria in human colorectal cancer. Gut pathogens, 6, 26.

35. Geng, J., Fan, H., Tang, X., Zhai, H. and Zhang, Z. (2013) Diversified pattern of the human colorectal cancer microbiome. Gut pathogens, 5, 2.

36. Wu, N., Yang, X., Zhang, R., Li, J., Xiao, X., Hu, Y., Chen, Y., Yang, F., Lu, N. and Wang, Z. (2013) Dysbiosis signature of fecal microbiota in colorectal cancer patients. Microbial ecology, 66, 462–470.

37. Wirbel, J., Pyl, P.T., Kartal, E., Zych, K., Kashani, A., Milanese, A., Fleck, J.S., Voigt, A.Y., Palleja, A. and Ponnudurai, R. (2019) Meta-analysis of fecal metagenomes reveals global microbial signatures that are specific for colorectal cancer. Nature medicine, 1.

38. Jenq, R.R., Taur, Y., Devlin, S.M., Ponce, D.M., Goldberg, J.D., Ahr, K.F., Littmann, E.R., Ling, L., Gobourne, A.C., Miller, L.C. et al. (2015) Intestinal Blautia Is Associated with Reduced Death from Graft-versus-Host Disease. Biol Blood Marrow Transplant, 21, 1373–1383.

39. Cancer Genome Atlas, N. (2012) Comprehensive molecular characterization of human colon and rectal cancer. Nature, 487, 330–337.

40. Langille, M.G., Zaneveld, J., Caporaso, J.G., McDonald, D., Knights, D., Reyes, J.A., Clemente, J.C., Burkepile, D.E., Thurber, R.L.V. and Knight, R. (2013) Predictive functional profiling of microbial communities using 16S rRNA marker gene sequences. Nature biotechnology, 31, 814.

41. Zeller, G., Tap, J., Voigt, A.Y., Sunagawa, S., Kultima, J.R., Costea, P.I., Amiot, A., Bohm, J., Brunetti, F., Habermann, N. et al. (2014) Potential of fecal microbiota for early-stage detection of colorectal cancer. Molecular Systems Biology, 10, 766–766.

42. Feng, Q., Liang, S., Jia, H., Stadlmayr, A., Tang, L., Lan, Z., Zhang, D., Xia, H., Xu, X., Jie, Z. et al. (2015) Gut microbiome development along the colorectal adenoma–carcinoma sequence. Nature Communications, 6.

