## Supplemental Materials for "Characterizing the Cancer-Associated Microbiome with Small RNA Sequencing Data"

***Corresponding author**

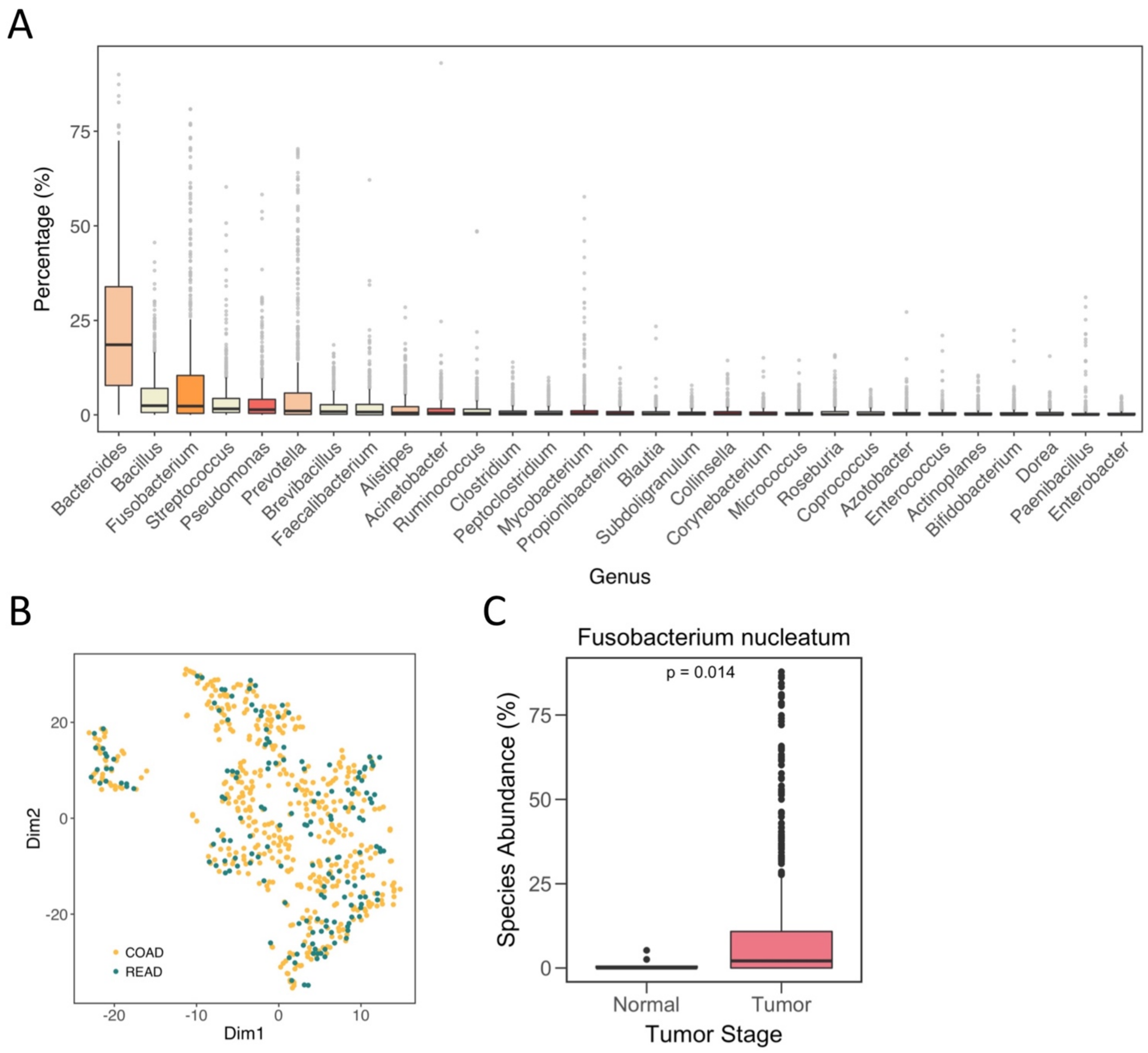


**Supplementary Figure S1. Phylogenetic profiles of the human gut microbiome in colorectal cancer.**

(A) Box plot showing the 30 most abundant genera in colorectal cancer. The order of genera is sorted according to the median of abundance. (B) Profiling of two types of cancer, COAD and READ, using two dimensions of t-SNE. (C) Relative abundance of *Fusobacterium nucleatum* in normal and tumor samples. Statistical significance was determined using the Mann–Whitney U test.

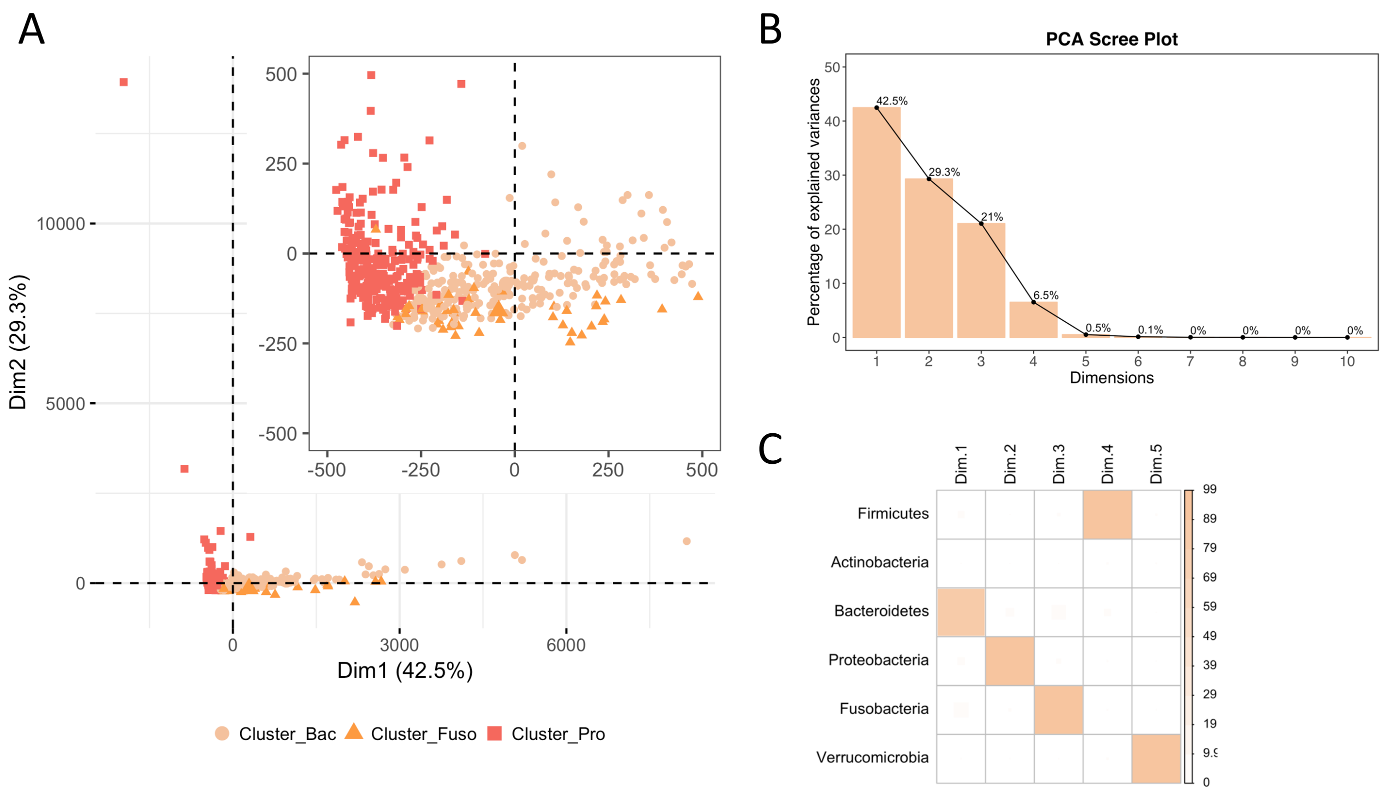


**Supplementary Figure S2. Principle component analysis (PCA) and PCA dimensions.**

(A) Principle component analysis showing separation of two major clusters, Cluster_Bac and Cluster_Pro. Colors and shapes indicate different clusters. (B) PCA screen plots explaining the variance of each dimension. (C) Correlation plots indicating correlation between phyla and PCA dimensions.

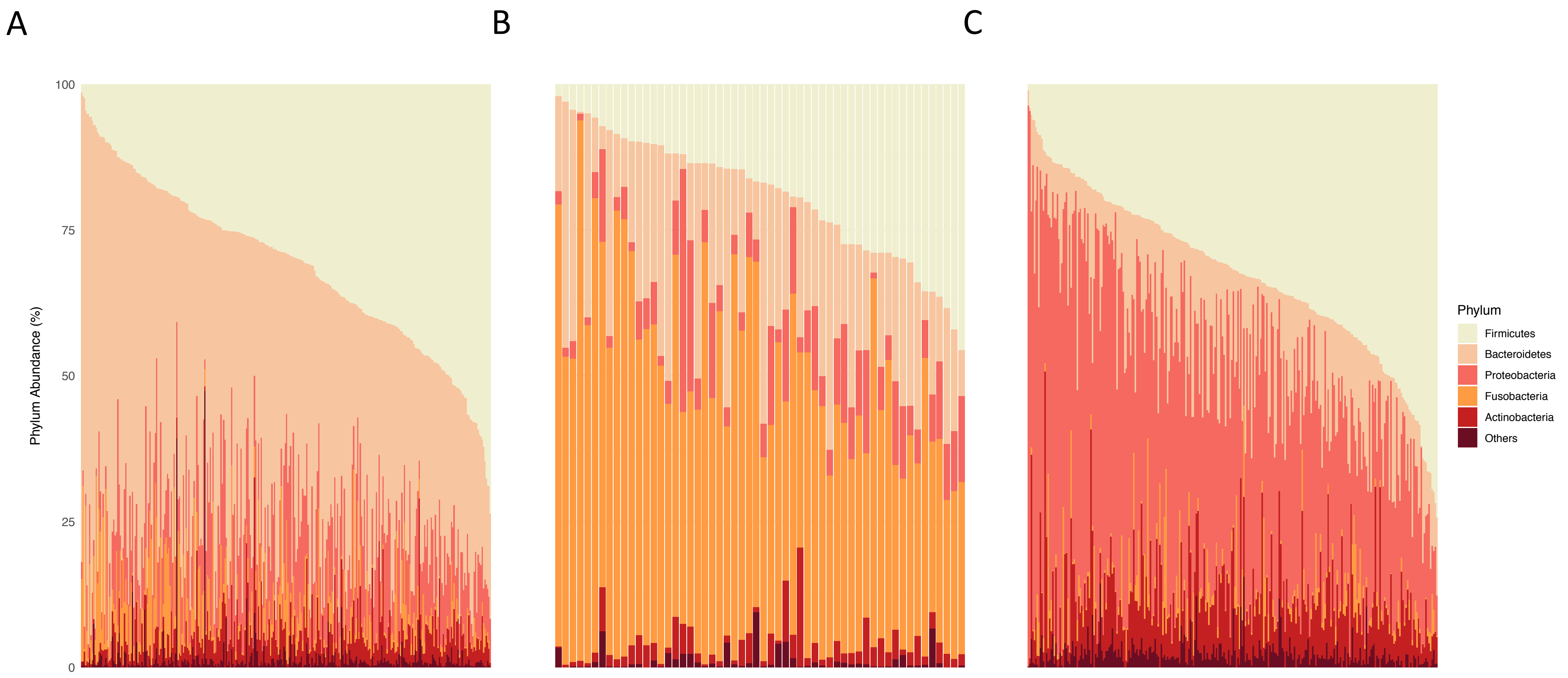


**Supplementary Figure S3. Stacked bar plot of phylum abundance among clusters.**

Three clusters, namely (A) Cluster_Bac, (B) Cluster_Fuso, and (C) Cluster_Pro, were used to perform an analysis of the relative phylum abundance in the clusters. Colors indicate different phyla.

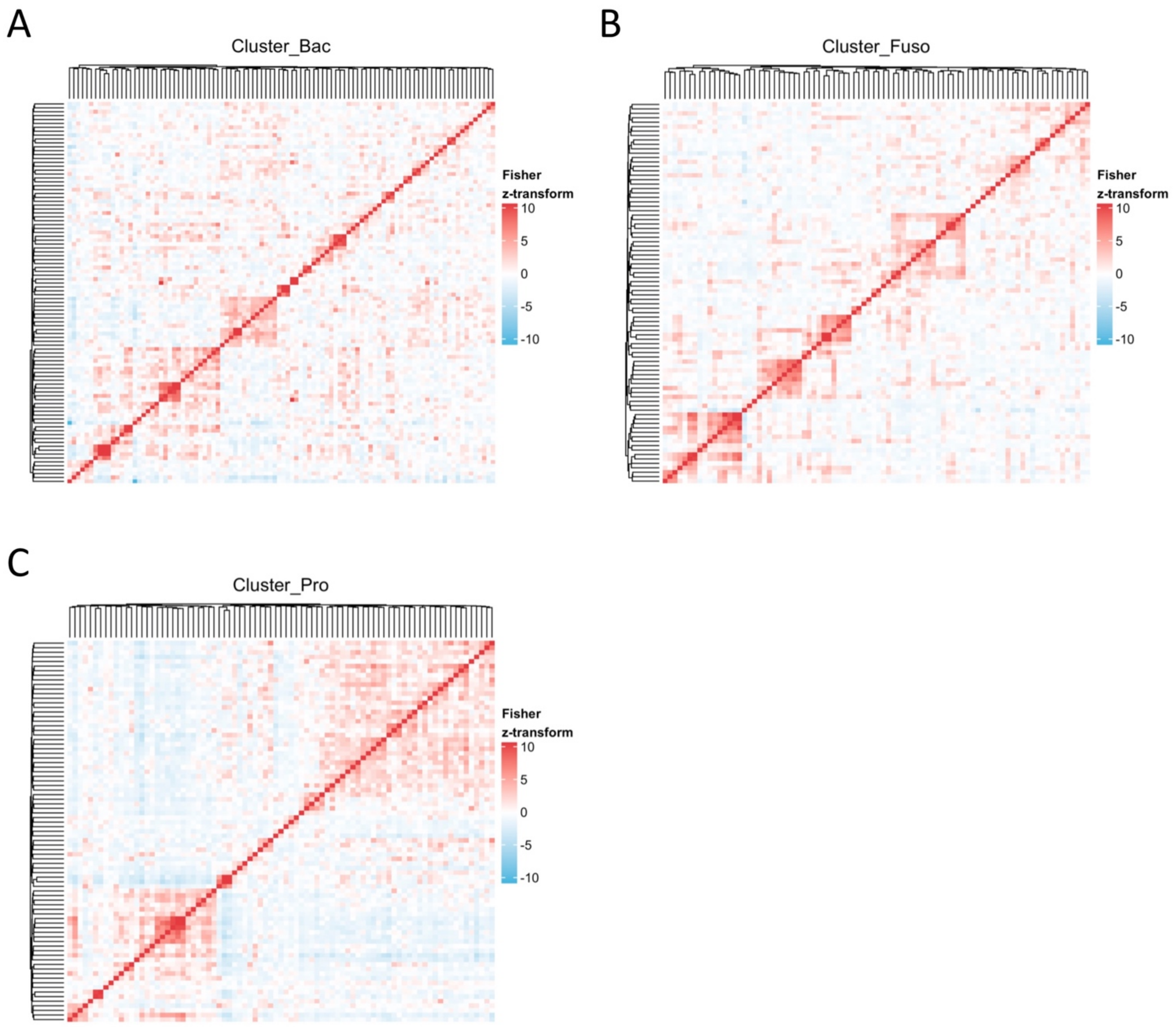


**Supplementary Figure S4. Bacterial co-occurrence heatmaps.**

(A–C) Co-occurrence heatmaps of bacterial genera in different clusters. Correlations were calculated using the Pearson correlation coefficient. Fisher z-transformation was used for normalization according to the sample size, rendering the networks comparable.

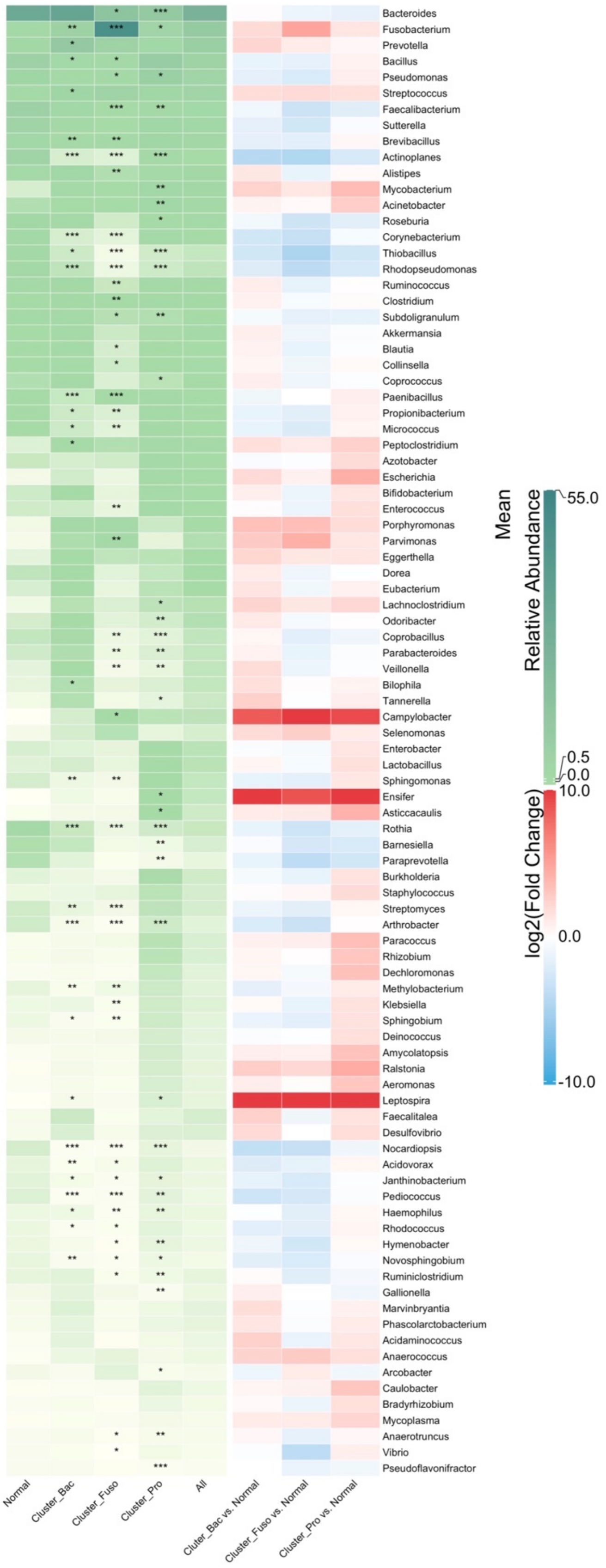


**Supplementary Figure S5. Common microbiome in CRC.**

This heatmap shows the common microbiome, which should be present in > 20% of the samples. The left heatmap (colored green) indicates the relative abundance of genera, and each column stands for three clusters (normal adjacent samples were removed), normal adjacent tissues, and all samples. The statistical significance of the three clusters was calculated between tumors in each cluster and all normal adjacent tissues using the Mann–Whitney U test (*$p \leq 0.05$ and $p > 0.01$; **$p \leq 0.01$ and $p > 0.001$; ***$p \leq 0.001$). The right heatmap colored with log2 (fold change). Fold change was defined as the genus relative abundance in clusters divided by the genus relative abundance in normal samples.

**Supplementary Table S1. Topological properties of co-occurrence network in Cluster_Bac**

| Genus | Average Shortest Path Length | Betweenness Centrality | Closeness Centrality | Clustering Coefficient | Degree | Neighborhood Connectivity | Radiality | Topological Coefficient |
|---|---|---|---|---|---|---|---|---|
| Acidaminococcus | 2.406 | 0.002 | 0.416 | 0.400 | 5 | 15.400 | 0.719 | 0.280 |
| Acidovorax | 2.365 | 0.002 | 0.423 | 0.472 | 9 | 16.778 | 0.727 | 0.311 |
| Acinetobacter | 2.156 | 0.018 | 0.464 | 0.305 | 15 | 15.333 | 0.769 | 0.231 |
| Actinoplanes | 2.156 | 0.021 | 0.464 | 0.325 | 16 | 14.313 | 0.769 | 0.216 |
| Adlercreutzia | 2.219 | 0.009 | 0.451 | 0.333 | 12 | 14.833 | 0.756 | 0.235 |
| Aeromonas | 2.531 | 0.005 | 0.395 | 0.286 | 7 | 12.286 | 0.694 | 0.276 |
| Akkermansia | 2.719 | 0.021 | 0.368 | 0.000 | 3 | 11.333 | 0.656 | 0.356 |
| Alistipes | 2.135 | 0.025 | 0.468 | 0.267 | 15 | 14.333 | 0.773 | 0.210 |
| Amycolatopsis | 2.188 | 0.015 | 0.457 | 0.244 | 13 | 14.154 | 0.763 | 0.210 |
| Anaerococcus | 2.688 | 0.003 | 0.372 | 0.000 | 4 | 8.750 | 0.663 | 0.287 |
| Anaerotruncus | 2.344 | 0.015 | 0.427 | 0.194 | 9 | 11.333 | 0.731 | 0.208 |
| Arcobacter | 2.531 | 0.002 | 0.395 | 0.333 | 4 | 13.750 | 0.694 | 0.314 |
| Arthrobacter | 2.010 | 0.022 | 0.497 | 0.385 | 22 | 16.045 | 0.798 | 0.216 |
| Asticcacaulis | 2.031 | 0.024 | 0.492 | 0.337 | 24 | 14.708 | 0.794 | 0.207 |
| Azotobacter | 2.750 | 0.000 | 0.364 | 0.500 | 5 | 14.200 | 0.650 | 0.418 |
| Bacillus | 1.927 | 0.025 | 0.519 | 0.310 | 19 | 16.737 | 0.815 | 0.197 |
| Bacteroides | 1.875 | 0.078 | 0.533 | 0.232 | 24 | 13.458 | 0.825 | 0.163 |
| Barnesiella | 2.583 | 0.002 | 0.387 | 0.300 | 5 | 12.400 | 0.683 | 0.305 |
| Bifidobacterium | 1.958 | 0.037 | 0.511 | 0.287 | 17 | 15.529 | 0.808 | 0.187 |
| Bilophila | 2.208 | 0.018 | 0.453 | 0.288 | 12 | 11.833 | 0.758 | 0.182 |
| Blautia | 2.406 | 0.003 | 0.416 | 0.286 | 7 | 15.857 | 0.719 | 0.294 |
| Bradyrhizobium | 2.531 | 0.005 | 0.395 | 0.286 | 7 | 13.714 | 0.694 | 0.295 |
| Brevibacillus | 1.906 | 0.029 | 0.525 | 0.255 | 18 | 15.667 | 0.819 | 0.180 |
| Burkholderia | 2.167 | 0.010 | 0.462 | 0.475 | 16 | 16.688 | 0.767 | 0.256 |
| Campylobacter | 2.552 | 0.005 | 0.392 | 0.200 | 6 | 9.833 | 0.690 | 0.232 |
| Caulobacter | 2.094 | 0.020 | 0.478 | 0.340 | 18 | 15.111 | 0.781 | 0.216 |
| Clostridium | 2.250 | 0.009 | 0.444 | 0.400 | 10 | 15.600 | 0.750 | 0.257 |

**Supplementary Table S1. Topological properties of co-occurrence network in Cluster_Bac (Continued)**

| Genus | Average Shortest Path Length | Betweenness Centrality | Closeness Centrality | Clustering Coefficient | Degree | Neighborhood Connectivity | Radiality | Topological Coefficient |
|---|---|---|---|---|---|---|---|---|
| Collinsella | 2.396 | 0.009 | 0.417 | 0.250 | 9 | 11.111 | 0.721 | 0.218 |
| Coprobacillus | 2.458 | 0.007 | 0.407 | 0.200 | 5 | 12.600 | 0.708 | 0.260 |
| Coprococcus | 2.063 | 0.021 | 0.485 | 0.368 | 17 | 15.412 | 0.788 | 0.213 |
| Corynebacterium | 1.979 | 0.028 | 0.505 | 0.352 | 23 | 15.739 | 0.804 | 0.204 |
| Dechloromonas | 2.146 | 0.010 | 0.466 | 0.407 | 14 | 17.929 | 0.771 | 0.263 |
| Deinococcus | 2.156 | 0.010 | 0.464 | 0.309 | 11 | 13.818 | 0.769 | 0.195 |
| Desulfotomaculum | 2.333 | 0.002 | 0.429 | 0.467 | 6 | 19.667 | 0.733 | 0.322 |
| Desulfovibrio | 2.406 | 0.009 | 0.416 | 0.250 | 9 | 10.778 | 0.719 | 0.218 |
| Dialister | 2.292 | 0.013 | 0.436 | 0.400 | 11 | 15.364 | 0.742 | 0.270 |
| Dorea | 2.010 | 0.012 | 0.497 | 0.450 | 19 | 16.316 | 0.798 | 0.215 |
| Eggerthella | 2.406 | 0.001 | 0.416 | 0.300 | 5 | 18.000 | 0.719 | 0.349 |
| Ensifer | 2.396 | 0.002 | 0.417 | 0.583 | 9 | 16.556 | 0.721 | 0.307 |
| Enterobacter | 1.979 | 0.023 | 0.505 | 0.379 | 20 | 16.800 | 0.804 | 0.215 |
| Enterococcus | 2.313 | 0.009 | 0.432 | 0.333 | 9 | 14.889 | 0.738 | 0.248 |
| Escherichia | 2.667 | 0.001 | 0.375 | 0.500 | 4 | 10.750 | 0.667 | 0.307 |
| Eubacterium | 2.021 | 0.015 | 0.495 | 0.458 | 18 | 16.167 | 0.796 | 0.215 |
| Faecalibacterium | 2.125 | 0.005 | 0.471 | 0.593 | 14 | 17.929 | 0.775 | 0.256 |
| Faecalitalea | 2.375 | 0.003 | 0.421 | 0.393 | 8 | 15.750 | 0.725 | 0.297 |
| Filifactor | 2.260 | 0.016 | 0.442 | 0.214 | 8 | 12.875 | 0.748 | 0.207 |
| Fusobacterium | 1.854 | 0.058 | 0.539 | 0.268 | 26 | 14.615 | 0.829 | 0.176 |
| Gallionella | 2.458 | 0.003 | 0.407 | 0.200 | 6 | 11.833 | 0.708 | 0.254 |
| Geobacillus | 2.208 | 0.004 | 0.453 | 0.422 | 10 | 17.300 | 0.758 | 0.253 |
| Holdemania | 2.208 | 0.013 | 0.453 | 0.291 | 11 | 14.545 | 0.758 | 0.220 |
| Hymenobacter | 2.333 | 0.007 | 0.429 | 0.289 | 10 | 13.800 | 0.733 | 0.245 |
| Janthinobacterium | 2.281 | 0.009 | 0.438 | 0.345 | 11 | 15.909 | 0.744 | 0.264 |

**Supplementary Table S1. Topological properties of co-occurrence network in Cluster_Bac (Continued)**

| Genus | Average Shortest Path Length | Betweenness Centrality | Closeness Centrality | Clustering Coefficient | Degree | Neighborhood Connectivity | Radiality | Topological Coefficient |
|---|---|---|---|---|---|---|---|---|
| Klebsiella | 2.615 | 0.003 | 0.382 | 0.200 | 5 | 9.400 | 0.677 | 0.250 |
| Lachnoclostridium | 2.438 | 0.013 | 0.410 | 0.190 | 7 | 10.000 | 0.713 | 0.207 |
| Lactobacillus | 2.885 | 0.000 | 0.347 | 1.000 | 2 | 13.000 | 0.623 | 0.619 |
| Leptospira | 2.333 | 0.003 | 0.429 | 0.545 | 11 | 17.545 | 0.733 | 0.308 |
| Marvinbryantia | 2.240 | 0.005 | 0.447 | 0.593 | 14 | 16.429 | 0.752 | 0.274 |
| Methylobacterium | 2.104 | 0.027 | 0.475 | 0.250 | 16 | 14.438 | 0.779 | 0.201 |
| Micrococcus | 1.979 | 0.027 | 0.505 | 0.338 | 21 | 15.714 | 0.804 | 0.201 |
| Mycobacterium | 2.208 | 0.008 | 0.453 | 0.306 | 9 | 16.222 | 0.758 | 0.242 |
| Mycoplasma | 2.281 | 0.002 | 0.438 | 0.524 | 7 | 17.143 | 0.744 | 0.268 |
| Odoribacter | 2.031 | 0.026 | 0.492 | 0.250 | 17 | 13.706 | 0.794 | 0.178 |
| Oscillibacter | 2.188 | 0.018 | 0.457 | 0.227 | 12 | 12.167 | 0.763 | 0.180 |
| Paenibacillus | 2.490 | 0.001 | 0.402 | 0.622 | 10 | 17.100 | 0.702 | 0.372 |
| Parabacteroides | 2.031 | 0.015 | 0.492 | 0.242 | 12 | 16.250 | 0.794 | 0.198 |
| Paracoccus | 2.219 | 0.016 | 0.451 | 0.314 | 15 | 15.267 | 0.756 | 0.239 |
| Paraprevotella | 2.135 | 0.017 | 0.468 | 0.133 | 10 | 12.300 | 0.773 | 0.171 |
| Parvimonas | 2.021 | 0.017 | 0.495 | 0.352 | 15 | 17.267 | 0.796 | 0.219 |
| Peptoclostridium | 2.052 | 0.022 | 0.487 | 0.292 | 16 | 14.375 | 0.790 | 0.189 |
| Phascolarctobacterium | 2.208 | 0.030 | 0.453 | 0.200 | 10 | 14.300 | 0.758 | 0.218 |
| Porphyromonas | 2.208 | 0.010 | 0.453 | 0.311 | 10 | 15.400 | 0.758 | 0.235 |
| Prevotella | 1.813 | 0.072 | 0.552 | 0.158 | 23 | 13.304 | 0.838 | 0.146 |
| Propionibacterium | 1.927 | 0.034 | 0.519 | 0.293 | 25 | 15.240 | 0.815 | 0.193 |
| Pseudoflavonifractor | 2.010 | 0.031 | 0.497 | 0.187 | 14 | 13.214 | 0.798 | 0.165 |
| Pseudomonas | 2.281 | 0.012 | 0.438 | 0.352 | 15 | 15.333 | 0.744 | 0.264 |
| Ralstonia | 2.156 | 0.012 | 0.464 | 0.341 | 14 | 15.786 | 0.769 | 0.229 |
| Rhizobium | 2.271 | 0.007 | 0.440 | 0.379 | 12 | 15.500 | 0.746 | 0.257 |

**Supplementary Table S1. Topological properties of co-occurrence network in Cluster_Bac (Continued)**

| Genus | Average Shortest Path Length | Betweenness Centrality | Closeness Centrality | Clustering Coefficient | Degree | Neighborhood Connectivity | Radiality | Topological Coefficient |
|---|---|---|---|---|---|---|---|---|
| Rhodococcus | 2.438 | 0.001 | 0.410 | 0.583 | 9 | 17.444 | 0.713 | 0.335 |
| Rhodopseudomonas | 2.219 | 0.010 | 0.451 | 0.327 | 11 | 16.091 | 0.756 | 0.251 |
| Roseburia | 2.063 | 0.010 | 0.485 | 0.510 | 18 | 15.944 | 0.788 | 0.221 |
| Rothia | 3.708 | 0.000 | 0.270 | 0.000 | 1 | 3.000 | 0.458 | 0.000 |
| Ruminiclostridium | 2.083 | 0.015 | 0.480 | 0.429 | 15 | 16.267 | 0.783 | 0.223 |
| Ruminococcus | 2.052 | 0.019 | 0.487 | 0.407 | 22 | 14.545 | 0.790 | 0.208 |
| Selenomonas | 3.135 | 0.000 | 0.319 | 1.000 | 2 | 7.500 | 0.573 | 0.682 |
| Sinorhizobium | 2.396 | 0.002 | 0.417 | 0.564 | 11 | 16.636 | 0.721 | 0.314 |
| Sphingobium | 2.010 | 0.030 | 0.497 | 0.319 | 21 | 15.238 | 0.798 | 0.205 |
| Sphingomonas | 2.260 | 0.009 | 0.442 | 0.408 | 16 | 15.438 | 0.748 | 0.262 |
| Staphylococcus | 2.656 | 0.002 | 0.376 | 0.500 | 5 | 11.200 | 0.669 | 0.333 |
| Streptococcus | 2.385 | 0.004 | 0.419 | 0.429 | 7 | 16.286 | 0.723 | 0.307 |
| Streptomyces | 2.146 | 0.015 | 0.466 | 0.381 | 15 | 15.000 | 0.771 | 0.221 |
| Subdoligranulum | 1.969 | 0.027 | 0.508 | 0.336 | 23 | 14.348 | 0.806 | 0.186 |
| Sutterella | 3.198 | 0.000 | 0.313 | 0.000 | 1 | 10.000 | 0.560 | 0.000 |
| Tannerella | 2.958 | 0.000 | 0.338 | 1.000 | 2 | 13.500 | 0.608 | 0.614 |
| Thiobacillus | 2.344 | 0.017 | 0.427 | 0.389 | 9 | 15.000 | 0.731 | 0.274 |
| Treponema | 2.354 | 0.012 | 0.425 | 0.286 | 7 | 14.286 | 0.729 | 0.257 |
| Veillonella | 3.042 | 0.000 | 0.329 | 0.000 | 2 | 8.000 | 0.592 | 0.500 |
| Vibrio | 2.292 | 0.006 | 0.436 | 0.289 | 10 | 14.400 | 0.742 | 0.240 |

**Supplementary Table S2. Topological properties of co-occurrence network in Cluster_Fuso**

| Genus | Average Shortest Path Length | Betweenness Centrality | Closeness Centrality | Clustering Coefficient | Degree | Neighborhood Connectivity | Radiality | Topological Coefficient |
|---|---|---|---|---|---|---|---|---|
| Acidovorax | 2.524 | 0.008 | 0.396 | 0.549 | 14 | 12.071 | 0.746 | 0.326 |
| Acinetobacter | 2.286 | 0.043 | 0.438 | 0.319 | 21 | 11.714 | 0.786 | 0.254 |
| Actinoplanes | 2.262 | 0.015 | 0.442 | 0.561 | 12 | 13.250 | 0.790 | 0.241 |
| Aeromonas | 2.429 | 0.014 | 0.412 | 0.357 | 8 | 9.375 | 0.762 | 0.215 |
| Akkermansia | 3.143 | 0.000 | 0.318 | 1.000 | 3 | 11.000 | 0.643 | 0.579 |
| Alistipes | 2.143 | 0.044 | 0.467 | 0.358 | 16 | 12.500 | 0.810 | 0.211 |
| Amycolatopsis | 2.357 | 0.017 | 0.424 | 0.286 | 8 | 9.625 | 0.774 | 0.201 |
| Anaerococcus | 3.226 | 0.000 | 0.310 | 0.000 | 2 | 6.500 | 0.629 | 0.550 |
| Arcobacter | 2.393 | 0.004 | 0.418 | 0.655 | 11 | 14.091 | 0.768 | 0.294 |
| Arthrobacter | 2.536 | 0.003 | 0.394 | 0.500 | 4 | 15.250 | 0.744 | 0.357 |
| Asticcacaulis | 2.571 | 0.007 | 0.389 | 0.607 | 8 | 13.000 | 0.738 | 0.348 |
| Azotobacter | 2.321 | 0.019 | 0.431 | 0.311 | 10 | 12.700 | 0.780 | 0.238 |
| Bacillus | 2.381 | 0.009 | 0.420 | 0.545 | 12 | 12.833 | 0.770 | 0.267 |
| Bacteroides | 2.869 | 0.005 | 0.349 | 0.000 | 3 | 8.667 | 0.688 | 0.348 |
| Barnesiella | 2.393 | 0.050 | 0.418 | 0.267 | 10 | 10.100 | 0.768 | 0.230 |
| Bifidobacterium | 2.226 | 0.027 | 0.449 | 0.333 | 13 | 13.000 | 0.796 | 0.232 |
| Bilophila | 2.655 | 0.010 | 0.377 | 0.524 | 7 | 10.000 | 0.724 | 0.365 |
| Blautia | 2.202 | 0.038 | 0.454 | 0.314 | 15 | 12.333 | 0.800 | 0.224 |
| Brachybacterium | 3.071 | 0.000 | 0.326 | 1.000 | 3 | 7.667 | 0.655 | 0.511 |
| Brevibacillus | 2.345 | 0.008 | 0.426 | 0.561 | 12 | 13.667 | 0.776 | 0.268 |
| Burkholderia | 2.345 | 0.026 | 0.426 | 0.560 | 14 | 13.214 | 0.776 | 0.281 |
| Campylobacter | 2.452 | 0.023 | 0.408 | 0.333 | 7 | 10.571 | 0.758 | 0.243 |
| Clostridium | 2.095 | 0.062 | 0.477 | 0.250 | 16 | 9.875 | 0.817 | 0.162 |
| Collinsella | 4.226 | 0.000 | 0.237 | 0.000 | 1 | 2.000 | 0.462 | 0.000 |
| Coprobacillus | 2.345 | 0.037 | 0.426 | 0.309 | 11 | 8.909 | 0.776 | 0.194 |
| Coprococcus | 2.238 | 0.031 | 0.447 | 0.404 | 17 | 12.000 | 0.794 | 0.226 |
| Corynebacterium | 2.238 | 0.032 | 0.447 | 0.399 | 18 | 12.556 | 0.794 | 0.251 |

**Supplementary Table S2. Topological properties of co-occurrence network in Cluster_Fuso (Continued)**

| Genus | Average Shortest Path Length | Betweenness Centrality | Closeness Centrality | Clustering Coefficient | Degree | Neighborhood Connectivity | Radiality | Topological Coefficient |
|---|---|---|---|---|---|---|---|---|
| Deinococcus | 2.274 | 0.021 | 0.440 | 0.470 | 12 | 12.667 | 0.788 | 0.226 |
| Desulfovibrio | 3.214 | 0.000 | 0.311 | 1.000 | 2 | 7.000 | 0.631 | 0.583 |
| Dialister | 3.179 | 0.001 | 0.315 | 0.000 | 2 | 7.000 | 0.637 | 0.545 |
| Dorea | 2.131 | 0.033 | 0.469 | 0.434 | 17 | 12.235 | 0.812 | 0.211 |
| Eggerthella | 2.536 | 0.005 | 0.394 | 0.429 | 7 | 10.286 | 0.744 | 0.286 |
| Ensifer | 2.643 | 0.007 | 0.378 | 0.400 | 6 | 9.167 | 0.726 | 0.296 |
| Enterobacter | 2.560 | 0.007 | 0.391 | 0.491 | 11 | 13.000 | 0.740 | 0.325 |
| Enterococcus | 2.548 | 0.009 | 0.393 | 0.464 | 8 | 11.500 | 0.742 | 0.280 |
| Eubacterium | 2.274 | 0.056 | 0.440 | 0.357 | 8 | 11.250 | 0.788 | 0.210 |
| Faecalibacterium | 2.036 | 0.070 | 0.491 | 0.304 | 19 | 12.211 | 0.827 | 0.194 |
| Faecalitalea | 2.262 | 0.015 | 0.442 | 0.491 | 11 | 12.818 | 0.790 | 0.237 |
| Filifactor | 3.107 | 0.006 | 0.322 | 0.000 | 2 | 7.500 | 0.649 | 0.500 |
| Frankia | 2.476 | 0.007 | 0.404 | 0.600 | 11 | 12.636 | 0.754 | 0.275 |
| Fusobacterium | 2.155 | 0.088 | 0.464 | 0.205 | 13 | 10.077 | 0.808 | 0.174 |
| Gallionella | 2.512 | 0.005 | 0.398 | 0.578 | 10 | 12.600 | 0.748 | 0.332 |
| Janthinobacterium | 2.250 | 0.034 | 0.444 | 0.487 | 13 | 12.538 | 0.792 | 0.241 |
| Lachnoclostridium | 2.214 | 0.058 | 0.452 | 0.227 | 12 | 8.833 | 0.798 | 0.164 |
| Lactobacillus | 2.500 | 0.025 | 0.400 | 0.509 | 11 | 11.000 | 0.750 | 0.250 |
| Leptospira | 2.381 | 0.035 | 0.420 | 0.306 | 9 | 11.111 | 0.770 | 0.239 |
| Leptotrichia | 2.917 | 0.000 | 0.343 | 1.000 | 2 | 12.000 | 0.681 | 0.600 |
| Marvinbryantia | 2.333 | 0.010 | 0.429 | 0.527 | 11 | 13.091 | 0.778 | 0.273 |
| Methylobacterium | 2.464 | 0.007 | 0.406 | 0.626 | 14 | 13.214 | 0.756 | 0.322 |
| Micrococcus | 2.310 | 0.033 | 0.433 | 0.367 | 16 | 12.750 | 0.782 | 0.255 |
| Mycobacterium | 2.821 | 0.001 | 0.354 | 0.600 | 5 | 11.800 | 0.696 | 0.407 |
| Mycoplasma | 2.274 | 0.022 | 0.440 | 0.333 | 9 | 11.778 | 0.788 | 0.214 |

**Supplementary Table S2. Topological properties of co-occurrence network in Cluster_Fuso (Continued)**

| Genus | Average Shortest Path Length | Betweenness Centrality | Closeness Centrality | Clustering Coefficient | Degree | Neighborhood Connectivity | Radiality | Topological Coefficient |
|---|---|---|---|---|---|---|---|---|
| Novosphingobium | 2.357 | 0.008 | 0.424 | 0.679 | 13 | 14.000 | 0.774 | 0.298 |
| Odoribacter | 2.429 | 0.015 | 0.412 | 0.556 | 10 | 11.900 | 0.762 | 0.277 |
| Paenibacillus | 2.393 | 0.005 | 0.418 | 0.727 | 11 | 13.818 | 0.768 | 0.300 |
| Parabacteroides | 2.333 | 0.018 | 0.429 | 0.500 | 12 | 12.667 | 0.778 | 0.264 |
| Paracoccus | 2.571 | 0.004 | 0.389 | 0.750 | 9 | 12.000 | 0.738 | 0.300 |
| Paraprevotella | 2.738 | 0.010 | 0.365 | 0.333 | 6 | 10.167 | 0.710 | 0.318 |
| Parvimonas | 2.857 | 0.003 | 0.350 | 0.333 | 4 | 9.000 | 0.690 | 0.350 |
| Pediococcus | 2.798 | 0.000 | 0.357 | 0.667 | 4 | 12.750 | 0.700 | 0.455 |
| Peptoclostridium | 2.179 | 0.049 | 0.459 | 0.407 | 14 | 12.071 | 0.804 | 0.214 |
| Phascolarctobacterium | 2.262 | 0.049 | 0.442 | 0.291 | 11 | 9.909 | 0.790 | 0.189 |
| Porphyromonas | 3.298 | 0.000 | 0.303 | 1.000 | 2 | 8.500 | 0.617 | 0.607 |
| Prevotella | 2.726 | 0.028 | 0.367 | 0.133 | 6 | 6.333 | 0.712 | 0.214 |
| Propionibacterium | 2.607 | 0.009 | 0.384 | 0.527 | 11 | 11.545 | 0.732 | 0.312 |
| Pseudoflavonifractor | 3.024 | 0.001 | 0.331 | 0.667 | 4 | 7.750 | 0.663 | 0.388 |
| Pseudomonas | 2.536 | 0.004 | 0.394 | 0.689 | 10 | 12.800 | 0.744 | 0.298 |
| Rhizobium | 2.369 | 0.010 | 0.422 | 0.571 | 14 | 13.571 | 0.772 | 0.302 |
| Rhodococcus | 2.321 | 0.016 | 0.431 | 0.491 | 11 | 12.364 | 0.780 | 0.242 |
| Rhodopseudomonas | 2.262 | 0.013 | 0.442 | 0.487 | 13 | 12.462 | 0.790 | 0.244 |
| Roseburia | 2.107 | 0.059 | 0.475 | 0.297 | 14 | 12.071 | 0.815 | 0.198 |
| Rothia | 2.500 | 0.004 | 0.400 | 0.778 | 9 | 12.778 | 0.750 | 0.290 |
| Ruminiclostridium | 2.512 | 0.021 | 0.398 | 0.345 | 11 | 9.182 | 0.748 | 0.242 |
| Ruminococcus | 2.393 | 0.004 | 0.418 | 0.667 | 10 | 14.000 | 0.768 | 0.311 |
| Selenomonas | 2.393 | 0.031 | 0.418 | 0.357 | 8 | 9.625 | 0.768 | 0.214 |
| Sphingomonas | 2.905 | 0.000 | 0.344 | 0.867 | 6 | 14.000 | 0.683 | 0.438 |
| Staphylococcus | 2.286 | 0.025 | 0.438 | 0.408 | 16 | 12.438 | 0.786 | 0.235 |

**Supplementary Table S2. Topological properties of co-occurrence network in Cluster_Fuso (Continued)**

| Genus | Average Shortest Path Length | Betweenness Centrality | Closeness Centrality | Clustering Coefficient | Degree | Neighborhood Connectivity | Radiality | Topological Coefficient |
|---|---|---|---|---|---|---|---|---|
| Streptococcus | 3.238 | 0.024 | 0.309 | 0.000 | 2 | 4.500 | 0.627 | 0.500 |
| Streptomyces | 2.393 | 0.006 | 0.418 | 0.712 | 12 | 14.167 | 0.768 | 0.315 |
| Subdoligranulum | 2.250 | 0.018 | 0.444 | 0.485 | 12 | 13.917 | 0.792 | 0.263 |
| Sutterella | 2.286 | 0.032 | 0.438 | 0.372 | 13 | 11.692 | 0.786 | 0.229 |
| Tannerella | 2.583 | 0.005 | 0.387 | 0.167 | 4 | 12.000 | 0.736 | 0.329 |
| Thiobacillus | 2.917 | 0.010 | 0.343 | 0.238 | 7 | 6.857 | 0.681 | 0.269 |
| Treponema | 3.667 | 0.001 | 0.273 | 0.000 | 2 | 4.000 | 0.556 | 0.500 |
| Veillonella | 2.750 | 0.008 | 0.364 | 0.467 | 6 | 7.833 | 0.708 | 0.290 |

**Supplementary Table S3. Topological properties of co-occurrence network in Cluster_Pro**

| Genus | Average Shortest Path Length | Betweenness Centrality | Closeness Centrality | Clustering Coefficient | Degree | Neighborhood Connectivity | Radiality | Topological Coefficient |
|---|---|---|---|---|---|---|---|---|
| Acidovorax | 2.304 | 0.000 | 0.434 | 0.667 | 7 | 20.857 | 0.783 | 0.409 |
| Acinetobacter | 2.468 | 0.000 | 0.405 | 0.000 | 1 | 44.000 | 0.755 | 0.000 |
| Actinoplanes | 2.304 | 0.006 | 0.434 | 0.214 | 8 | 12.875 | 0.783 | 0.260 |
| Aeromonas | 2.063 | 0.010 | 0.485 | 0.327 | 11 | 17.273 | 0.823 | 0.262 |
| Agrobacterium | 2.139 | 0.008 | 0.467 | 0.455 | 12 | 17.833 | 0.810 | 0.317 |
| Akkermansia | 2.076 | 0.005 | 0.482 | 0.306 | 9 | 21.111 | 0.821 | 0.318 |
| Alistipes | 1.937 | 0.017 | 0.516 | 0.505 | 20 | 17.400 | 0.844 | 0.260 |
| Amycolatopsis | 1.899 | 0.012 | 0.527 | 0.341 | 14 | 18.714 | 0.850 | 0.253 |
| Arthrobacter | 1.924 | 0.021 | 0.520 | 0.359 | 18 | 17.389 | 0.846 | 0.256 |
| Asticcacaulis | 1.608 | 0.064 | 0.622 | 0.327 | 36 | 17.639 | 0.899 | 0.235 |
| Azotobacter | 2.380 | 0.000 | 0.420 | 1.000 | 3 | 28.667 | 0.770 | 0.585 |
| Bacillus | 1.848 | 0.029 | 0.541 | 0.407 | 23 | 17.174 | 0.859 | 0.245 |
| Bacteroides | 1.481 | 0.181 | 0.675 | 0.238 | 44 | 15.955 | 0.920 | 0.210 |
| Bdellovibrio | 2.266 | 0.001 | 0.441 | 0.711 | 10 | 20.300 | 0.789 | 0.398 |
| Bifidobacterium | 2.253 | 0.012 | 0.444 | 0.236 | 11 | 12.636 | 0.791 | 0.248 |
| Bilophila | 2.114 | 0.003 | 0.473 | 0.578 | 10 | 19.700 | 0.814 | 0.313 |
| Blautia | 1.785 | 0.044 | 0.560 | 0.372 | 26 | 18.192 | 0.869 | 0.252 |
| Bradyrhizobium | 2.139 | 0.002 | 0.467 | 0.491 | 11 | 20.364 | 0.810 | 0.339 |
| Brevibacillus | 1.835 | 0.032 | 0.545 | 0.374 | 20 | 18.850 | 0.861 | 0.258 |
| Brevundimonas | 2.203 | 0.005 | 0.454 | 0.467 | 10 | 18.500 | 0.800 | 0.330 |
| Burkholderia | 1.873 | 0.020 | 0.534 | 0.476 | 22 | 18.591 | 0.854 | 0.273 |
| Caulobacter | 2.013 | 0.007 | 0.497 | 0.451 | 18 | 18.722 | 0.831 | 0.301 |
| Clostridium | 2.038 | 0.007 | 0.491 | 0.509 | 11 | 19.545 | 0.827 | 0.287 |
| Collinsella | 2.557 | 0.000 | 0.391 | 0.700 | 5 | 18.600 | 0.741 | 0.503 |
| Coprobacillus | 2.304 | 0.004 | 0.434 | 0.533 | 6 | 16.667 | 0.783 | 0.324 |
| Coprococcus | 1.899 | 0.017 | 0.527 | 0.458 | 18 | 19.167 | 0.850 | 0.269 |
| Corynebacterium | 1.835 | 0.018 | 0.545 | 0.424 | 21 | 19.476 | 0.861 | 0.271 |

**Supplementary Table S3. Topological properties of co-occurrence network in Cluster_Pro (Continued)**

| Genus | Average Shortest Path Length | Betweenness Centrality | Closeness Centrality | Clustering Coefficient | Degree | Neighborhood Connectivity | Radiality | Topological Coefficient |
|---|---|---|---|---|---|---|---|---|
| Dechloromonas | 1.975 | 0.014 | 0.506 | 0.395 | 20 | 17.650 | 0.838 | 0.276 |
| Deinococcus | 2.051 | 0.012 | 0.488 | 0.364 | 11 | 16.455 | 0.825 | 0.246 |
| Dorea | 1.810 | 0.028 | 0.552 | 0.435 | 24 | 18.667 | 0.865 | 0.256 |
| Eggerthella | 2.228 | 0.004 | 0.449 | 0.556 | 9 | 18.444 | 0.795 | 0.335 |
| Ensifer | 2.190 | 0.001 | 0.457 | 0.472 | 9 | 20.111 | 0.802 | 0.347 |
| Enterobacter | 2.342 | 0.001 | 0.427 | 0.267 | 6 | 17.667 | 0.776 | 0.369 |
| Enterococcus | 2.215 | 0.004 | 0.451 | 0.429 | 7 | 18.714 | 0.797 | 0.326 |
| Escherichia | 3.785 | 0.025 | 0.264 | 0.000 | 2 | 1.500 | 0.536 | 0.500 |
| Eubacterium | 1.861 | 0.012 | 0.537 | 0.542 | 20 | 20.850 | 0.857 | 0.286 |
| Faecalibacterium | 1.772 | 0.041 | 0.564 | 0.387 | 27 | 18.333 | 0.871 | 0.251 |
| Faecalitalea | 2.354 | 0.002 | 0.425 | 0.489 | 10 | 16.900 | 0.774 | 0.352 |
| Flavobacterium | 2.114 | 0.010 | 0.473 | 0.321 | 8 | 14.125 | 0.814 | 0.219 |
| Fusobacterium | 2.076 | 0.007 | 0.482 | 0.250 | 9 | 18.444 | 0.821 | 0.292 |
| Gemmata | 1.911 | 0.011 | 0.523 | 0.510 | 18 | 19.611 | 0.848 | 0.284 |
| Hyphomicrobium | 1.962 | 0.009 | 0.510 | 0.390 | 15 | 19.133 | 0.840 | 0.273 |
| Janthinobacterium | 2.570 | 0.002 | 0.389 | 0.300 | 5 | 12.000 | 0.738 | 0.347 |
| Klebsiella | 2.823 | 0.050 | 0.354 | 0.000 | 2 | 10.000 | 0.696 | 0.500 |
| Lachnoclostridium | 2.671 | 0.000 | 0.374 | 0.700 | 5 | 11.800 | 0.722 | 0.393 |
| Lactobacillus | 2.772 | 0.000 | 0.361 | 0.000 | 2 | 11.000 | 0.705 | 0.500 |
| Leptospira | 1.937 | 0.019 | 0.516 | 0.384 | 20 | 18.200 | 0.844 | 0.272 |
| Leptothrix | 2.114 | 0.006 | 0.473 | 0.485 | 17 | 17.882 | 0.814 | 0.319 |
| Methylobacterium | 1.987 | 0.009 | 0.503 | 0.425 | 18 | 17.667 | 0.835 | 0.272 |
| Micrococcus | 1.772 | 0.024 | 0.564 | 0.395 | 24 | 19.625 | 0.871 | 0.265 |
| Mycobacterium | 2.203 | 0.003 | 0.454 | 0.600 | 5 | 24.800 | 0.800 | 0.407 |
| Nocardiopsis | 1.886 | 0.091 | 0.530 | 0.327 | 18 | 18.111 | 0.852 | 0.262 |

**Supplementary Table S3. Topological properties of co-occurrence network in Cluster_Pro (Continued)**

| Genus | Average Shortest Path Length | Betweenness Centrality | Closeness Centrality | Clustering Coefficient | Degree | Neighborhood Connectivity | Radiality | Topological Coefficient |
|---|---|---|---|---|---|---|---|---|
| Novosphingobium | 2.051 | 0.012 | 0.488 | 0.372 | 13 | 17.692 | 0.825 | 0.275 |
| Odoribacter | 2.418 | 0.003 | 0.414 | 0.472 | 9 | 14.222 | 0.764 | 0.323 |
| Paenibacillus | 2.570 | 0.002 | 0.389 | 0.400 | 5 | 11.200 | 0.738 | 0.306 |
| Parabacteroides | 2.076 | 0.003 | 0.482 | 0.576 | 12 | 19.917 | 0.821 | 0.311 |
| Paracoccus | 2.051 | 0.022 | 0.488 | 0.255 | 11 | 15.455 | 0.825 | 0.235 |
| Pediococcus | 2.101 | 0.005 | 0.476 | 0.533 | 15 | 19.200 | 0.816 | 0.337 |
| Peptoclostridium | 2.620 | 0.002 | 0.382 | 0.333 | 4 | 12.500 | 0.730 | 0.371 |
| Propionibacterium | 1.722 | 0.046 | 0.581 | 0.316 | 27 | 18.037 | 0.880 | 0.240 |
| Pseudomonas | 1.924 | 0.022 | 0.520 | 0.353 | 18 | 17.389 | 0.846 | 0.251 |
| Ralstonia | 2.063 | 0.016 | 0.485 | 0.418 | 11 | 18.182 | 0.823 | 0.287 |
| Rhizobium | 1.823 | 0.029 | 0.549 | 0.373 | 25 | 18.200 | 0.863 | 0.256 |
| Rhodococcus | 1.924 | 0.014 | 0.520 | 0.340 | 18 | 19.000 | 0.846 | 0.271 |
| Rhodopseudomonas | 2.089 | 0.007 | 0.479 | 0.628 | 13 | 19.231 | 0.819 | 0.310 |
| Roseburia | 1.899 | 0.010 | 0.527 | 0.561 | 19 | 20.211 | 0.850 | 0.285 |
| Rothia | 2.671 | 0.000 | 0.374 | 1.000 | 3 | 17.000 | 0.722 | 0.607 |
| Ruminococcus | 2.367 | 0.002 | 0.422 | 0.636 | 11 | 19.182 | 0.772 | 0.417 |
| Shewanella | 2.633 | 0.000 | 0.380 | 0.667 | 4 | 12.000 | 0.728 | 0.387 |
| Sphingobium | 1.899 | 0.016 | 0.527 | 0.437 | 20 | 18.800 | 0.850 | 0.280 |
| Sphingomonas | 1.911 | 0.011 | 0.523 | 0.509 | 19 | 20.579 | 0.848 | 0.294 |
| Sphingopyxis | 2.013 | 0.011 | 0.497 | 0.417 | 16 | 17.938 | 0.831 | 0.276 |
| Staphylococcus | 2.089 | 0.007 | 0.479 | 0.278 | 9 | 15.000 | 0.819 | 0.238 |
| Streptococcus | 2.215 | 0.004 | 0.451 | 0.536 | 8 | 18.125 | 0.797 | 0.318 |
| Streptomyces | 2.038 | 0.014 | 0.491 | 0.352 | 15 | 17.333 | 0.827 | 0.271 |
| Subdoligranulum | 1.861 | 0.015 | 0.537 | 0.516 | 20 | 20.450 | 0.857 | 0.280 |
| Sutterella | 1.962 | 0.007 | 0.510 | 0.429 | 15 | 22.467 | 0.840 | 0.321 |

**Supplementary Table S3. Topological properties of co-occurrence network in Cluster_Pro (Continued)**

| Genus | Average Shortest Path Length | Betweenness Centrality | Closeness Centrality | Clustering Coefficient | Degree | Neighborhood Connectivity | Radiality | Topological Coefficient |
|---|---|---|---|---|---|---|---|---|
| Thiobacillus | 2.532 | 0.001 | 0.395 | 0.667 | 6 | 16.833 | 0.745 | 0.432 |
| Ureaplasma | 2.127 | 0.002 | 0.470 | 0.500 | 9 | 20.444 | 0.812 | 0.325 |
| Veillonella | 4.772 | 0.000 | 0.210 | 0.000 | 1 | 2.000 | 0.371 | 0.000 |
